# A modular framework for the development of targeted Covid-19 blood transcript profiling panels

**DOI:** 10.1101/2020.05.20.107243

**Authors:** Darawan Rinchai, Basirudeen Kabeer, Mohammed Toufiq, Zohreh Calderone, Sara Deola, Tobias Brummaier, Mathieu Garand, Ricardo Branco, Nicole Baldwin, Mohamed Alfaki, Matthew Altman, Alberto Ballestrero, Matteo Bassetti, Gabriele Zoppoli, Andrea De Maria, Benjamin Tang, Davide Bedognetti, Damien Chaussabel

## Abstract

Covid-19 morbidity and mortality are associated with a dysregulated immune response. Tools are needed to enhance existing immune profiling capabilities in affected patients. Here we aimed to develop an approach to support the design of focused blood transcriptome panels for profiling the immune response to SARS-CoV-2 infection. We designed a pool of candidates based on a pre-existing and well-characterized repertoire of blood transcriptional modules. Available Covid-19 blood transcriptome data was also used to guide this process. Further selection steps relied on expert curation. Additionally, we developed several custom web applications to support the evaluation of candidates. As a proof of principle, we designed three targeted blood transcript panels, each with a different translational connotation: therapeutic development relevance, SARS biology relevance and immunological relevance. Altogether the work presented here may contribute to the future expansion of immune profiling capabilities via targeted profiling of blood transcript abundance in Covid-19 patients.

## INTRODUCTION

Covid-19 is an infectious, respiratory disease caused by a newly discovered coronavirus: SARS-CoV-2. The severity of symptoms and the course of infection vary widely, with most patients presenting mild symptoms. However, about 20% of patients develop severe disease and require hospitalization (1,2). The interaction between innate and adaptive immunity can lead to the development of neutralizing antibodies against SARS-CoV-2 antigens that might be associated with viral clearance and protection (3). But immune factors are also believed to play an important role in the rapid clinical deterioration observed in some Covid-19 patients (4). There is thus a need to develop new modalities that can improve the delineation of “immune trajectories” during SARS-CoV-2 infection.

Blood transcriptome profiling involves measuring the abundance of circulating leukocyte RNA on a genome-wide scale (5). Processing of the samples and the raw sequencing data however, is time consuming and requires access to sophisticated laboratory and computational infrastructure. Thus, the possibility of implementing this approach on large scales to ensure immediate translational potential is limited. Such unbiased omics profiling data might rather be leveraged to inform the development of more practical, scalable and targeted transcriptional profiling assays. These assays could in turn serve to significantly bolster existing immune profiling capacity.

Fixed sets of transcripts grouped based on co-expression observed in large collections of reference datasets provide a robust platform for transcriptional profiling data analyses (6). Here we leveraged a repertoire of 382 transcriptional modules previously developed by our team (7). The repertoire is based on a collection of reference patient cohorts encompassing 16 pathological or physiological states and 985 individual transcriptome profiles. In this proof of principle study, we used the available transcript profiling data from two separate studies to select Covid-19 relevant sets of modules (8,9). Next, we applied filters based on pre-specified selection criteria (e.g. immunologic relevance or therapeutic relevance). Finally, expert curation was used as the last selection step. For this we have developed custom web applications to consolidate the information necessary for the evaluation of candidates. One of these applications provides access to module-level transcript abundance profiles for available Covid-19 blood transcriptome profiling datasets. Another web interface was implemented which serves as a scaffold for the juxtaposition of such transcriptional profiling data with extensive functional annotations.

## RESULTS

### Mapping Covid-19 blood transcriptome signatures against a pre-existing reference set of transcriptional modules

SARS-CoV-2 infection might not result in changes in transcript abundance across all of the 382 transcriptional modules constituting our repertoire [(encompassing 14,168 transcripts) see methods section and (7)]. Indeed, the 16 reference patient cohorts upon which our repertoire was based encompass a wide range of immune states, including infectious diseases but also autoimmune diseases, pregnancy and cancer. Thus, the first step involved identifying subsets of modules for which changes could be observed in Covid-19 patients.

We used two sets of Covid-19 patients for this proof of principle analysis. These datasets were contributed by Xiong *et al.* (9) (one control and three subjects) and Ong *et al.* (8) (nine controls and three subjects profiled at multiple time points). Their data were generated using RNA-seq and Nanostring technology, respectively. The generic 594 transcript panel used by Ong *et al.* did not give sufficient coverage across the 382-module set. We thus mapped the transcript changes at a lower resolution, using a framework formed by 38 module “aggregates”. These 38 aggregates were constituted by grouping modules based on similarities in abundance patterns across the 16 reference datasets [see methods section and (7)].

We first assessed changes in transcript abundance resulting from SARS-CoV-2 infection across the 38 module aggregates (**Figure 1**). In general, we saw a decrease in aggregates associated with lymphocytic compartments (aggregates A1 & A5) and an increase in aggregates associated with myeloid compartments and inflammation (aggregates A33 & A35). As expected, we also saw increases over uninfected controls for the module aggregate associated with interferon (IFN) responses (A28) and the module aggregate presumably associated with the effector humoral response (A27). We detected a wide spread of values for aggregate A11 for the Nanostring (Ong *et al.*) dataset. However, this aggregate comprises only one module, with only two of its transcripts measured in this Nanostring code set (the probe coverage across all aggregates is shown in **Supplementary Figure 1**).

**Figure 1:**
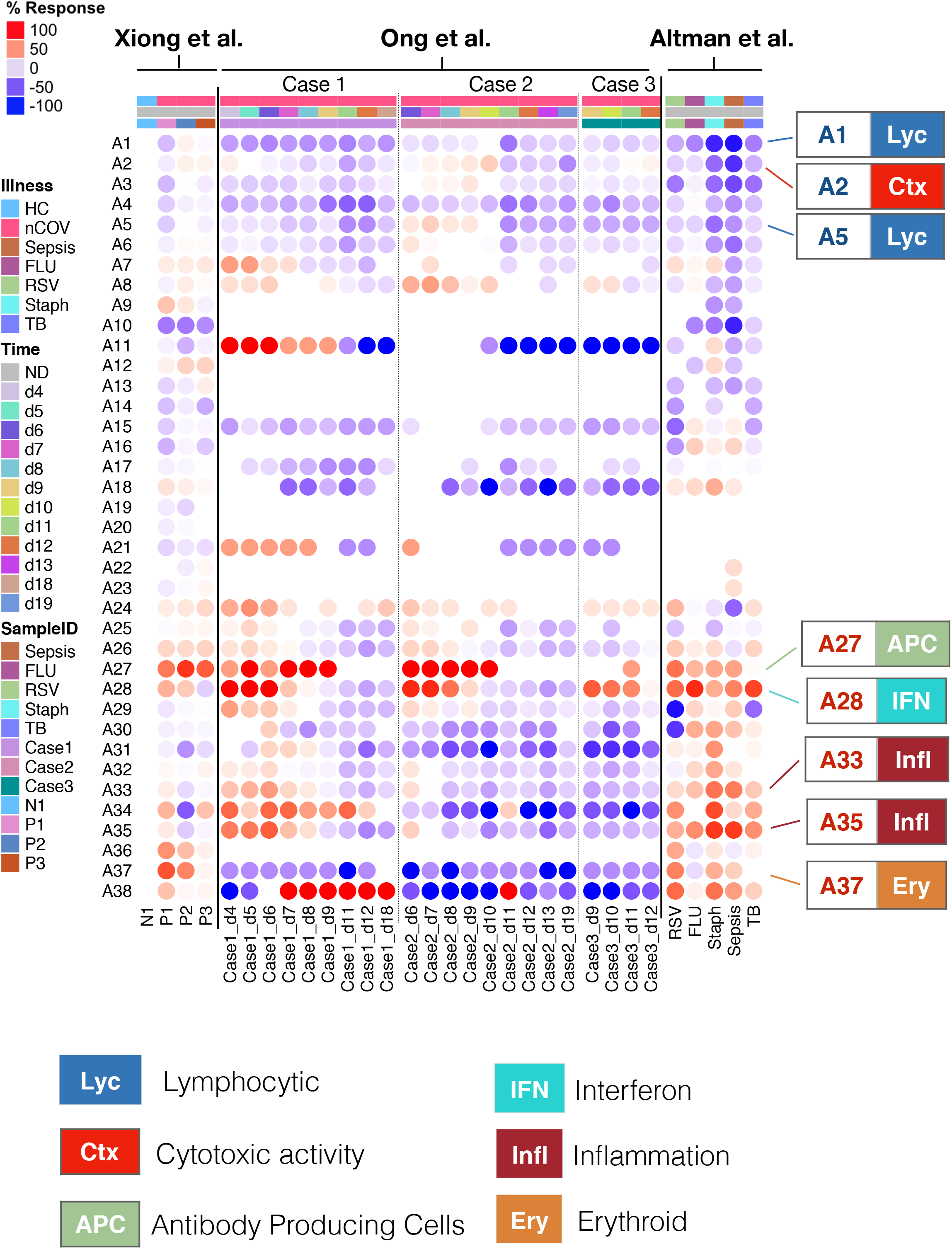
Mapping Covid-19 blood transcriptome signatures at the module aggregate level. The columns on this heatmap represent samples (Xiong *et al.* & Ong *et al.*) or patient cohorts (Altman *et al.*). Module aggregates (A1-A38) are arranged as rows. The colored spots represent the proportion of transcripts comprising each transcriptional module aggregate found to be differentially expressed compared to control samples. The cutoffs vary from one study to another due to differences in the design and the profiling platforms used. Thus, module aggregate response values range from 100% (all transcripts comprised in the module aggregate increased) to −100% (all decreased). The Xiong *et al.* dataset comprised one control and three Covid-19 patients and transcript abundance was measured by RNA-seq. The Ong *et al.* dataset comprised three Covid-19 cases from whom samples were collected serially, and nine uninfected controls (8). Transcript abundance was measured using a 594 gene standard immune panel from Nanostring. Patterns are also shown for cohorts comprised in the Altman *et al.* dataset (7). The colored labels (right) indicate functional associations for some of the aggregates.

Despite large differences between the two studies in terms of design, range of clinical severity, technology platforms and module coverage, the combined overall changes (detected at a high-level perspective) are consistent with those observed in known acute infections, such as those caused by influenza, respiratory syncytial virus (RSV) or *S. aureus.* This consistency is evidenced by the indicated by patterns of change observed for the reference fingerprints shown alongside those of Covid-19 patients (**Figure 1**).

Overall, such a high-level analysis allows us to identify module aggregates forming our repertoire that may be included in further selection steps. In this proof of principle analysis, we selected 17 aggregates for further analysis (**Table 1**).

**Table 1:** List of Covid-19 relevant aggregates and module sets

| Module Aggregate | Module Set | Modules | Functional annotations |
| --- | --- | --- | --- |
| **A1** | **A1/S1** | M14.42, M15.38, M12.6, M13.27, | T cells |
|  | **A1/S2** | M14.23, M15.87, M14.5, M14.49, M12.1, M14.20 | Gene transcription |
|  | **A1/S3** | M12.8, M15.29, M14.58, M15.51, M14.64, M16.78, M14.75, M15.82, M14.80 | B cells |
| **A2** | **A2/S1** | M13.21, M9.1 | Cytotoxic lymphocytes |
|  | **A2/S2** | M14.13, M14.72, M13.13, M13.14, M14.45, M13.10, M15.91 | TBD |
| **A4** | **A4/S1** | M16.69, M16.72, M16.50, M16.77 | Antigen presentation, |
| **A5** | **A5/S1** | M16.95, M16.36 | B cells |
|  | **A5/S2** | M16.57, M16.18, M16.65, M16.111, M16.99 | B cells |
| **A7** | **A7/S1** | M15.61 | Monocytes |
| **A8** | **A8/S1** | M16.30 | Complement |
|  | **A8/S2** | M16.106 | TBD |
| **A10** | **A10/S1** | M15.102 | Prostanoids |
| **A26** | **A26/S1** | M12.2 | Monocytes |
| **A27** | **A27/S1** | M13.32, M12.15, M16.92, M15.110, M16.60 | Antibody producing cells |
| **A28** | **A28/S1** | M15.127, M8.3 | Interferon response |
|  | **A28/S2** | M15.64 | Interferon response |
|  | **A28/S3** | M15.86, M10.1, M13.17 | Interferon response |
| **A31** | **A31/S1** | M16.64 | Platelet/Prostaglandin |
|  | **A31/S2** | M15.58 | Monocytes |
| **A33** | **A33/S1** | M15.104, M14.82, M14.24, M15.108 | Cytokines/chemokines, Inflammation |
|  | **A33/S2** | M14.19, M14.76, M14.50, M14.26, M16.101, M16.100, M16.80 | Inflammation |
| **A34** | **A34/S1** | M14.59, M10.3, M16.109, M8.2 | Platelets, Prostanoids |
| **A35** | **A35/S1** | M14.65, M14.28, M15.81, M16.79, M13.3, M14.7, | Monocytes, Neutrophils |
|  | **A35/S2** | M15.26, M12.10, M13.22, M15.109, M15.78, M13.16, | Neutrophils, Inflammation |
| **A36** | **A36/S1** | M16.34, M16.82, M15.97, M14.51, M15.118, M16.88 | Gene transcription |
| **A37** | **A37/S1** | M9.2, M14.53, M11.3, M12.11, M15.100 M15.74, M13.26, M13.30, M15.53, | Erythroid cells |
| **A38** | **A38/S1** | M10.4 | Neutrophil activation |
|  | **A38/S2** | M16.96, M12.9, M14.68 | Erythroid cells |

### Identification of coherent sets of Covid-19-relevant modules

The abundance patterns for modules comprised in a given aggregate are not always homogeneous (**Figure 2**). Thus, a next step would consist of identifying sets of modules within an aggregate that display coherent abundance patterns across modules forming a given aggregate.

**Figure 2.**
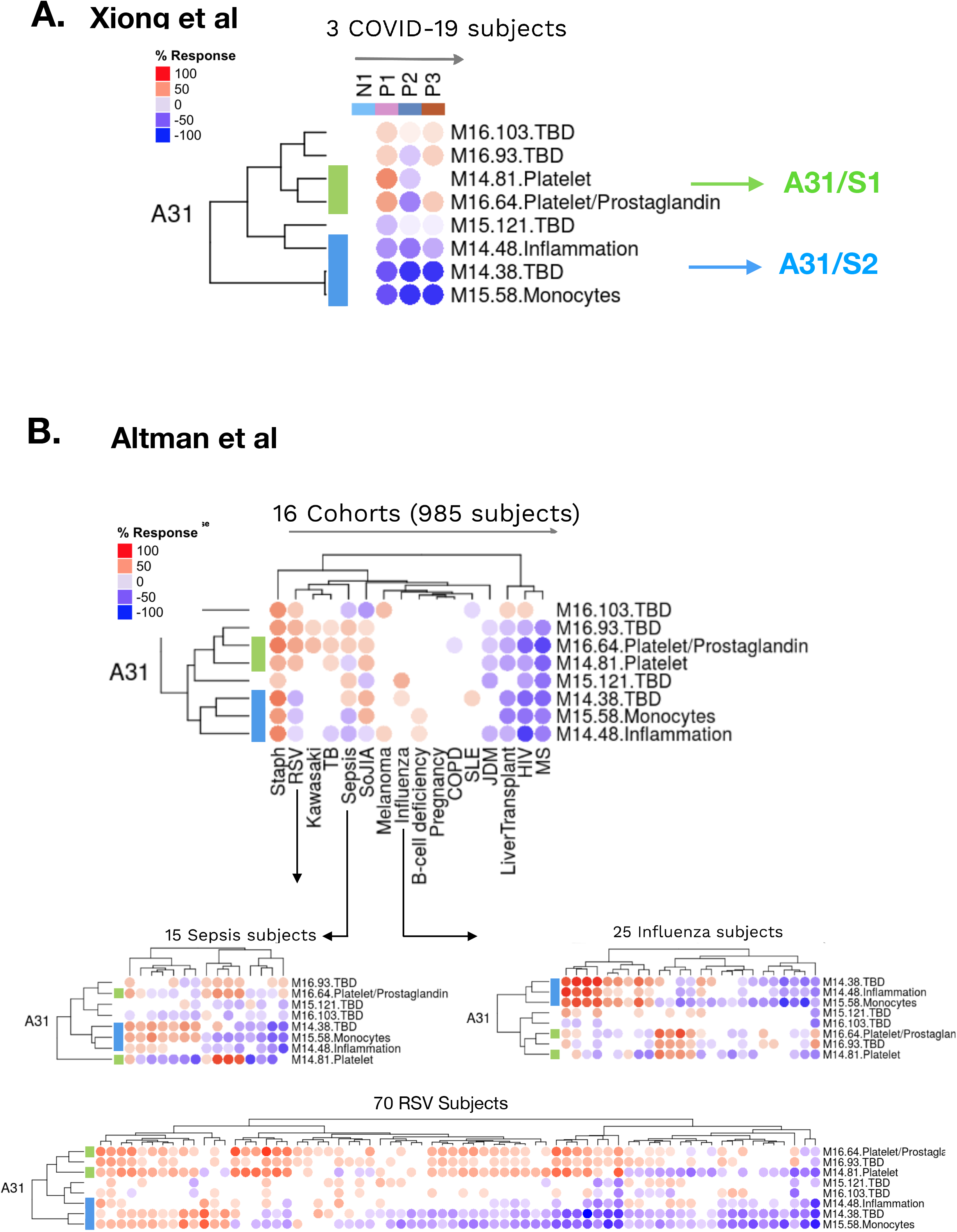
Delineation of sets of Covid-19 relevant A31 modules. A. Transcript abundance profiles of A31 modules in Covid-19 patients. This heatmap represents the abundance levels for transcripts forming modules belonging to aggregate A31 (rows), across three Covid-19 patients (P1-P3) relative to one uninfected control subject (columns). The data are expressed as the proportion of constitutive transcripts in each module being significantly increased (red circles) or decreased (blue circles) relative to N1. **B. Transcript abundance profiles of A31 modules in reference disease cohorts.** The top heatmap represents the abundance levels for transcripts forming modules belonging to aggregate A31 (rows), across 16 reference patient cohorts (columns). The bottom heatmaps represent the changes in abundance across the individuals comprised in two relevant patient cohorts, including pediatric patients with severe influenza or RSV infection and adult patients with sepsis.

To achieve this, we first mapped the changes in transcript abundance associated with Covid-19 disease using the RNAseq dataset from Xiong *et al*., as illustrated for A31 (**Figure 2A**) and A28 (**Figure 3A**). Similar plots can be generated for all other aggregates using the “COVID-19” web application (also compiled in **Supplementary File 1** and listed in **Table 2**). Next, we identified and assigned a module set ID for each aggregate the modules that formed homogeneous clusters. For example, we designated the first A28 set as A28/S1. Such module grouping is only based on patterns of transcript abundance observed in three Covid-19 patients; however, the groupings were often consistent with those observed for the much larger reference cohorts that constitute the module repertoire (**Figure 2B** and **Figure 3B**). A28/S1, which is formed by M8.3 and M15.127, serves as a good example of this consistency (**Figure 3B**). Likewise, the segregation of the modules forming A31 based on differences observed in the three Covid-19 patients was also apparent in the reference patient cohorts (**Figure 2B**). Specifically, an increase in A31/S1 modules, which accompanied a decrease in A31/S2 modules, in these three patients was also characteristic of RSV patients.

**Figure 3.**
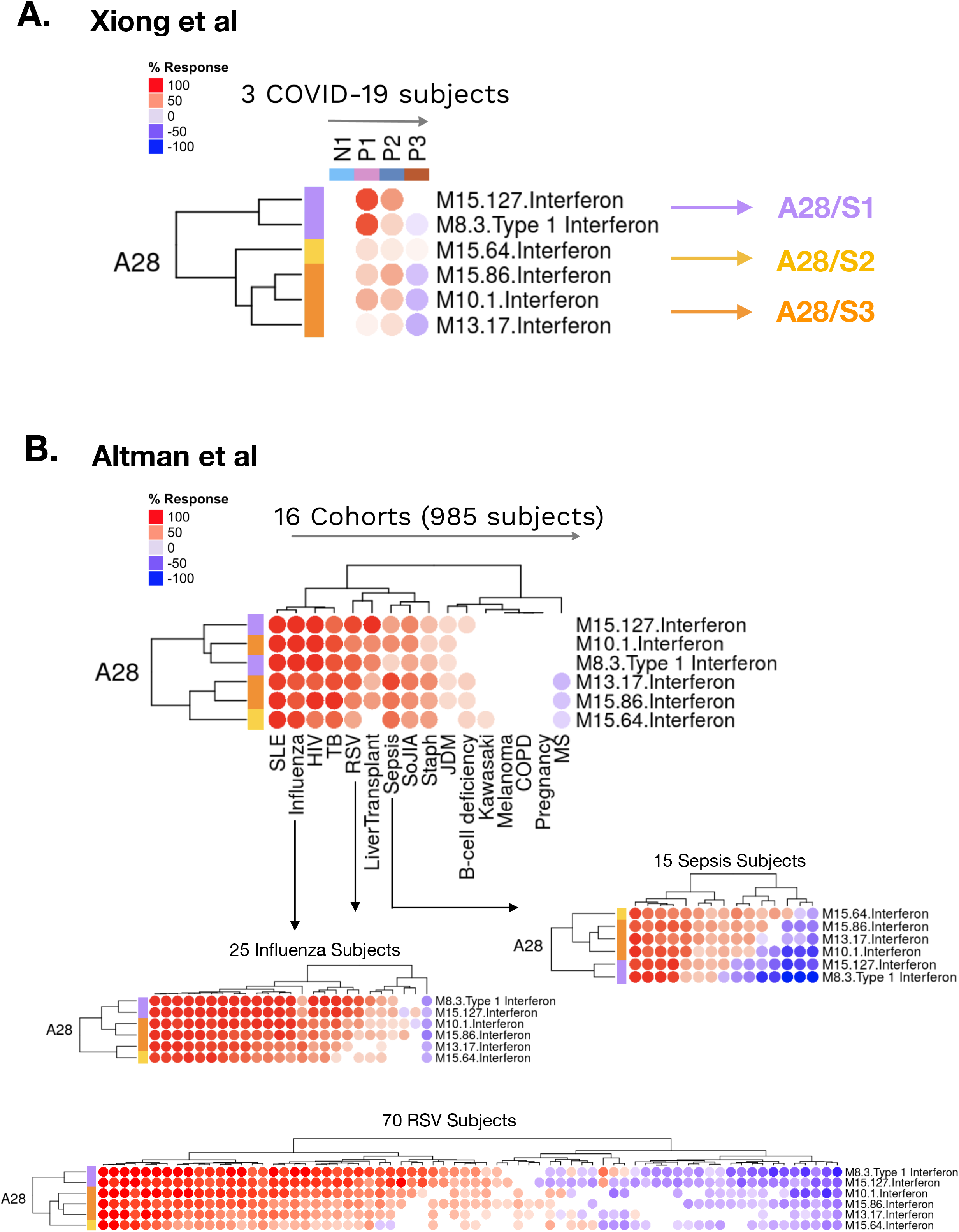
Delineation of sets of Covid-19 relevant A28 modules. A. Transcript abundance profiles of A28 modules in Covid-19 patients. This heatmap shows the abundance levels for transcripts forming modules belonging to aggregate A28 (rows), across three Covid-19 patients (P1-P3) relative to one uninfected control subject (columns). The data are expressed as the proportion of constitutive transcripts in each module being significantly increased (red circles) or decreased (blue circles) relative to N1 **B. Transcript abundance profiles of A28 modules in reference disease cohorts.** The top heatmap shows the abundance levels for transcripts forming modules belonging to aggregate A28 (rows), across 16 reference patient cohorts (columns). The bottom heatmaps show changes in abundance across individuals constituting the two relevant patient cohorts, including pediatric patients with severe influenza or RSV infection and adult patients with sepsis.

**Table 2:**
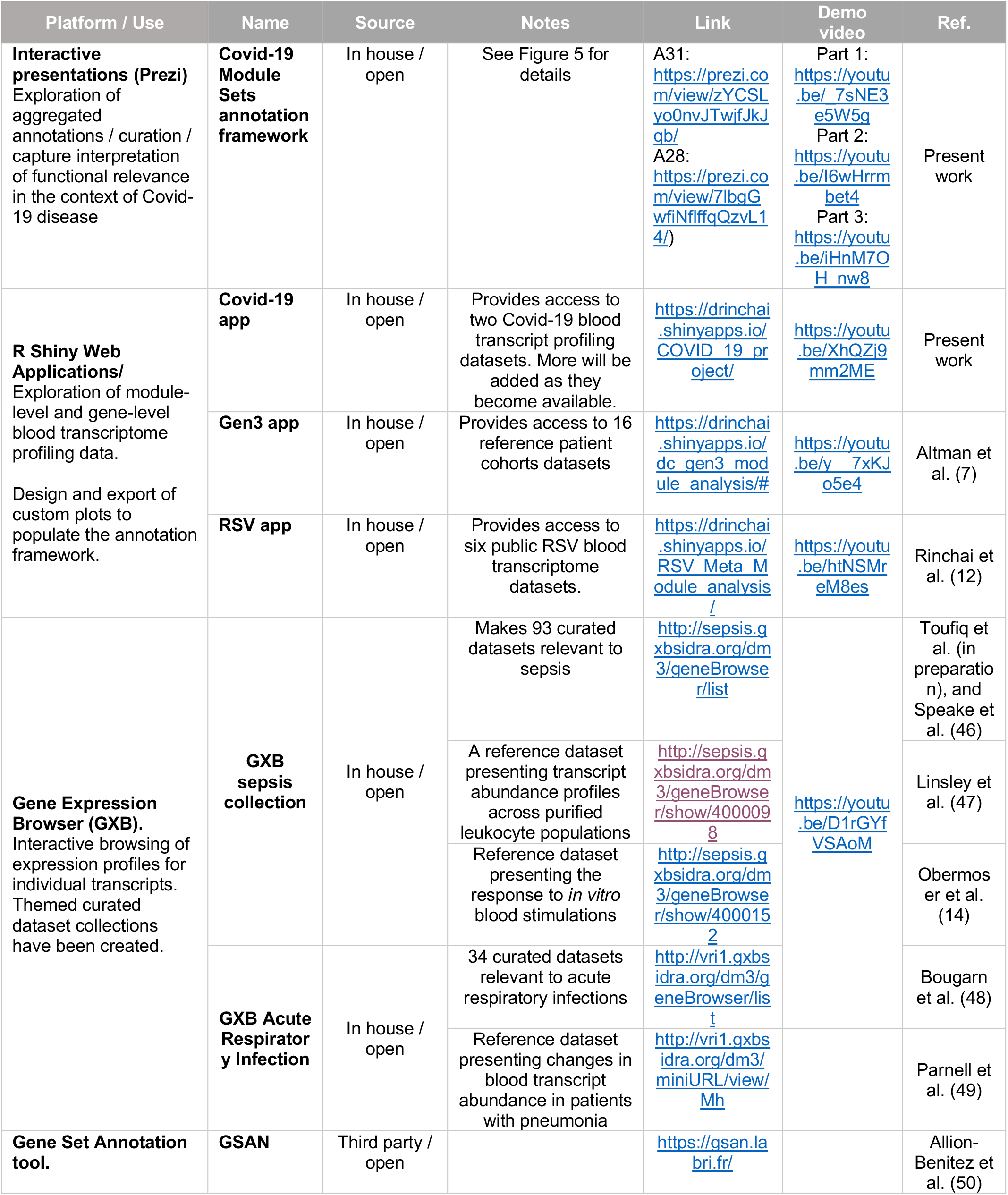
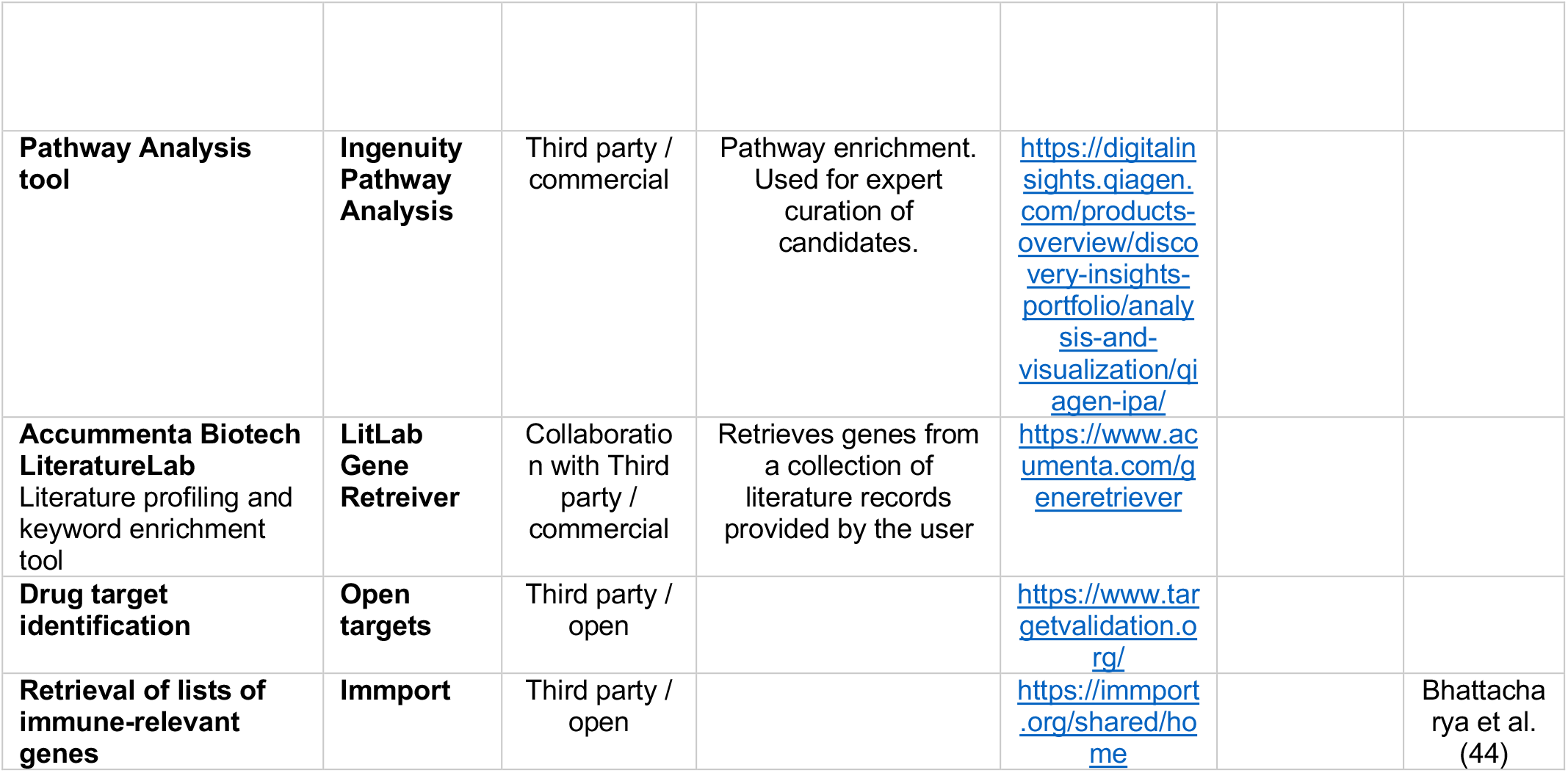
Resources used for annotation and interpretation

We ultimately derived 28 homogeneous Covid-19 relevant module sets from the 17 aggregates selected in the earlier step (**Table 1**). These sets might be used as a basis for further selection.

### Design of an illustrative targeted panel emphasizing immunological relevance

In the previous step, we used available Covid-19 data to guide the selection of 28 distinct “Covid-19 relevant module sets”. In the next step, we selected the transcripts within each module set that warranted inclusion in one of three illustrative Covid-19 targeted panels. A first panel was formed using immunologic relevance as the primary criterion, a second was formed on the basis of relevance to coronavirus biology, a third was constituted on the basis of relevance to therapy.

For the first panel we matched transcripts comprised in each module set to a list of canonical immune genes (see methods for details). Expert curation also involved accessing transcript profiling data from the reference datasets, indicating for instance leukocyte restriction or patterns of response to a wide range of immune stimuli *in vitro*. We describe our approach for module and gene annotation in more detail below and provide access to our resources to support expert curation (**Table 2**).

For our illustrative case, we selected one representative transcript per module set to produce a panel comprised of 28 representative transcripts (**Table 3)**. Examples of signatures surveyed by such a panel include: **1) ISG15** in A28/S1 (interferon responses), which encodes for a member of the ubiquitin family. ISG15 plays a central role in the host defense to viral infections (10). **2) GATA1** in A37/S1 (erythroid cells), which encodes for a master regulator of erythropoiesis (11). It is associated with a module signature (A37) that we recently reported as being associated with immunosuppressive states, such as late stage cancer and maintenance immunosuppressive therapy in solid organ transplant recipients (12). In the same report we also found an association between this signature and heightened severity in patients with RSV infection and established a putative link with a population of immunosuppressive circulating erythroid cells (13). **3) CD38** in A27/S1 (cell cycle), which encodes for the CD38 molecule expressed on different circulating leukocyte populations. In whole blood we find the abundance of its transcript to correlates with that of IGJ, TNFRSF17 (BCMA), TXNDC5 (M12.15). Such a signature was previously found to be increased in response to vaccination at day 7 post administration, to correlate with the prevalence of antibody producing cells, and the development of antibody titers at day 28 (14). **4) TLR8** in A35/S1 (inflammation), encodes toll-like receptor 8. Expression of transcripts comprising this aggregate is generally restricted to neutrophils and robustly increased during sepsis (e.g. as we have described in detail earlier for ACSL1, another transcript belonging to this aggregate (15)). **5) GZMB** in A2/S1 (Cytotoxic cells) encodes Granzyme B, a serine protease known to play a role in immune-mediated cytotoxicity. Other transcripts forming this panel are listed in **Table 3**.

**Table 3:**
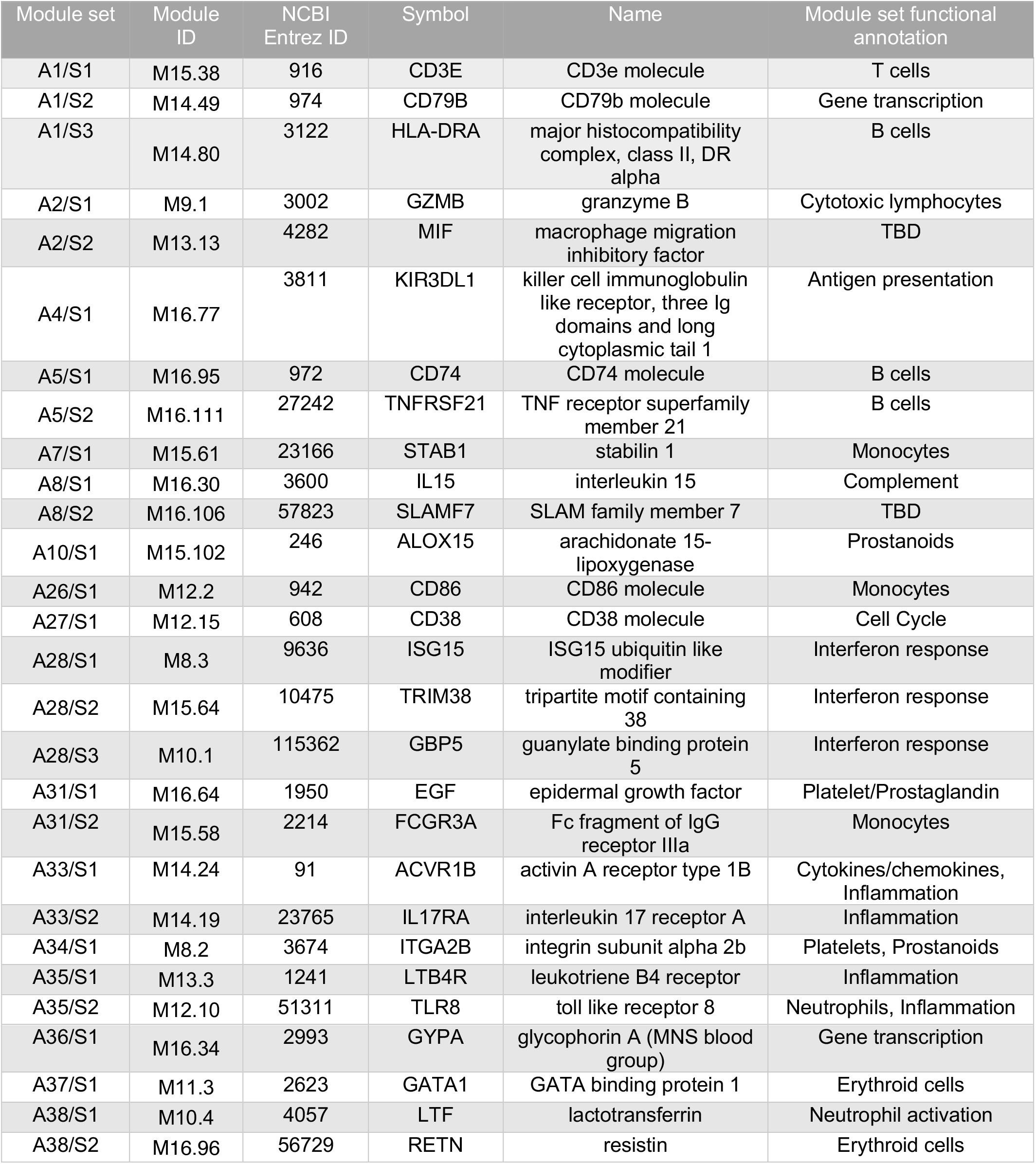
Illustrative targeted panel – Immunology relevance focus

Even with the limited amount of data available to guide the selection in the previous steps, it is reasonable to assume that such a panel (while not optimal) would already provide valid information for Covid-19 immune profiling. Additional Covid-19 blood transcriptome data that will become available in the coming weeks will allow us to refine the overall selection process.

### Design of an illustrative targeted panel emphasizing therapeutic relevance

A different translational connotation was given for this second panel. Here, we based the selection on the same collection of 28 module sets. However, this time, whenever possible, we included transcripts that could have value as targets for the treatment of Covid-19 patients. An initial screen identified 82 transcripts encoding molecules that are known targets for existing drugs (see Methods). We further prioritized these candidates based on an expert’s evaluation of the compatibility of use of the drugs for treating Covid-19 patients. As an exception, module sets belonging to A28 (interferon response) were selected based on their suitability as markers of a response to interferon therapy. Sets for which no targets of clinical relevance were identified (16/28) were instead represented in the panel by immunologically-relevant transcripts (defined earlier).

We ultimately identified a preliminary set of 12 targets through this high stringency selection process (**Table 4**). Developing effective immune modulation therapies in critical care settings has proven challenging (16). Current efforts in the context of Covid-19 disease particularly aim at controlling runaway systemic immune responses or so called “cytokine storms” that have been associated with organ damage and clinical worsening. Targets of interest identified among our gene set include: **1) IL6R** in A35/S2 (inflammation), encoding the Interleukin-6 Receptor, which is a target for the biologic drug Tocilizumab. Several studies have tested this antagonist in open label single arm trials in Covid-19 patients with the intent of blocking the cytokine storm associated with severe Covid-19 infection (17,18). **2) CCR2** in A26/1 (monocytes), encoding the chemokine (C-C motif) receptor 2, is targeted along with CCR5 by the drug Cenicriviroc. This drug exerts potent anti-inflammatory activity (19). **3) TBXA2R** in A31/1 (platelets), encoding the Thromboxane receptor, is targeted by several drugs with anti-platelet aggregation properties (20). **4) PDE8A** in A33/S1 (inflammation), encoding Phosphodiesterase 8A, is targeted by Pentoxifylline, a non-selective phosphodiesterase inhibitor that increases perfusion and may reduce risk of acute kidney injury and attenuates LPS-induced inflammation (21). **5) NQO1** in A8/S1 (Complement), encoding NAD(P)H quinone dehydrogenase 1. The NQO1 antagonist Vatiquinone (EPI-743) has been found to inhibit ferroptosis (22), a process associated with tissue injury (23), including in sepsis (24). A complete list is provided in **Table 4**.

**Table 4:**
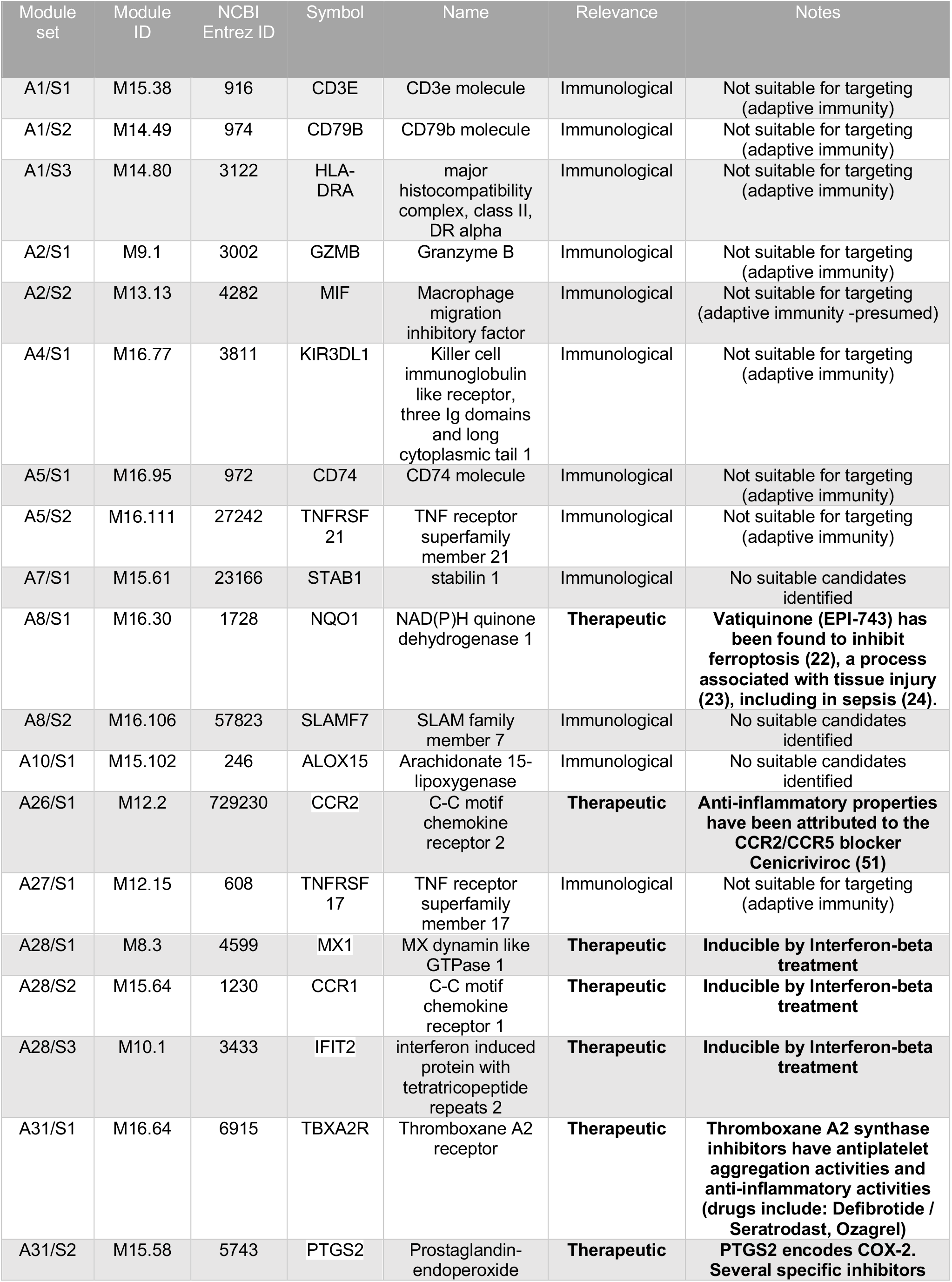
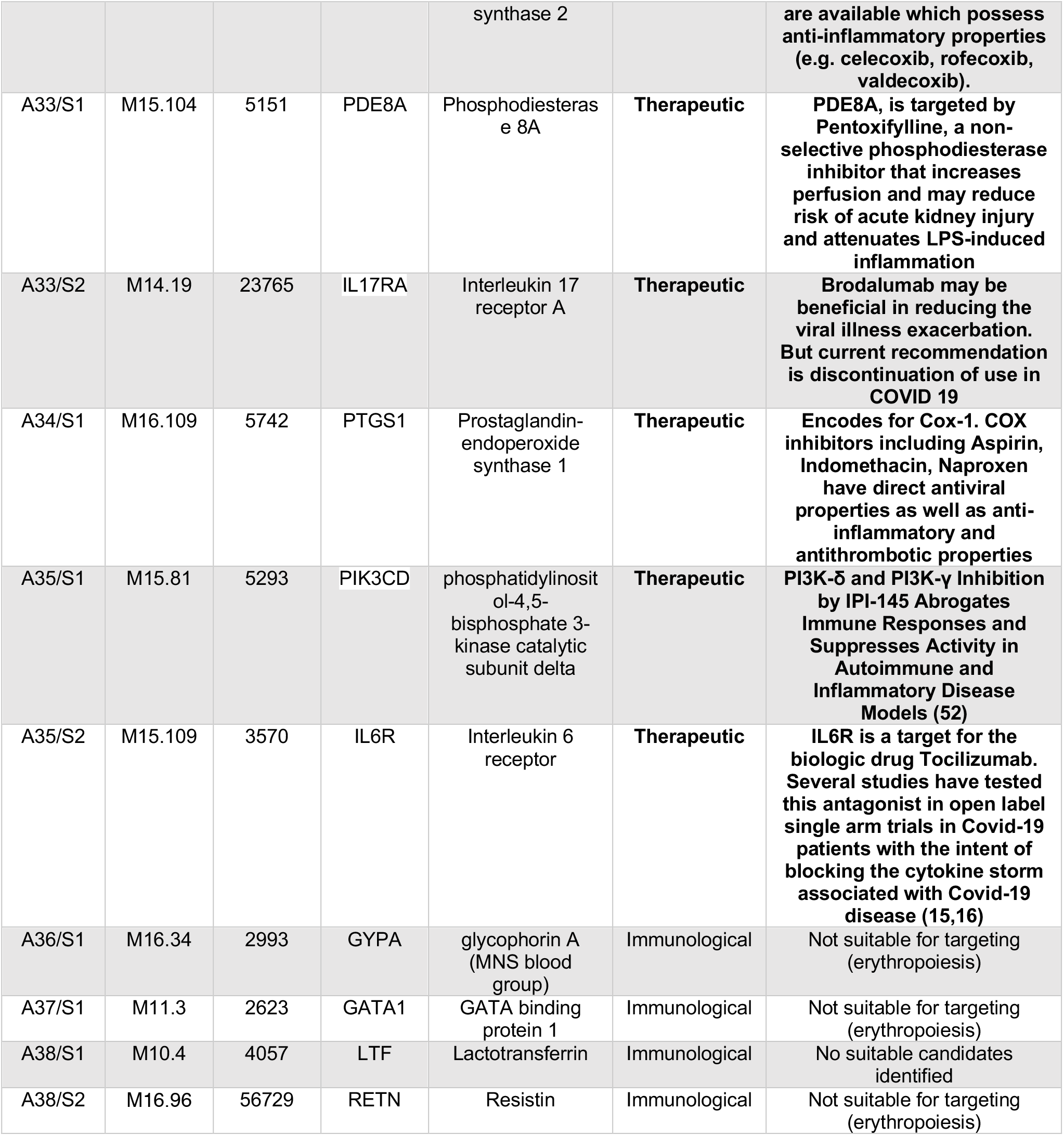
Illustrative targeted panel – Therapeutic relevance focus

The fact that this transcript panel and the previous survey the same pre-defined 28 homogenous Covid-19 relevant module sets should make them largely synonymous (since modules are formed on the basis of co-expression). Nevertheless, this second panel may be more relevant for investigators interested in investigating new therapeutic approaches or measuring responses to treatment.

### Design of a targeted panel of blood transcripts of relevance for SARS-CoV-2 biology

For the third panel designed in this proof of principle, we primarily selected transcripts based on their relevance to SARS biology. As a first step, we used a literature profiling tool to identify SARS, MERS, or Covid-19 literature articles that were associated with transcripts forming the 28 Covid-19 module sets. Next, the potential associations were subjected to expert curation (see Methods). Once again, to keep redundancies to a minimum, we only included one candidate per set in this panel (**Table 5**). Notable examples include: **1) LTF** in A38/S1 (neutrophil activation) encodes Lactotransferrin, that is known to block the binding of the SARS-CoV spike protein to host cells, thus exerting an inhibitory function at the viral attachment stage (25). **2) FURIN** in A37/S1 (Erythroid cells), encodes a proprotein convertase that preactivates SARS-CoV-2, thus reducing its dependence on target cell proteases for entry (26). **3) EGR1** in A7/S1 (Monocytes), encodes Early Growth Response 1, which upon induction by SARS Coronavirus Papain-Like Protease mediates up-Regulation of TGF-β1 (27). **4) STAT1** in A28/S3 (Interferon response), encodes a transcription factor known to play an important role in the induction of antiviral effector responses. It was reported that SARS ORF6 Antagonizes STAT1 function by preventing its translocation to the nucleus and acts as an interferon antagonist in the context of SARS-CoV infection (28).

**Table 5:**
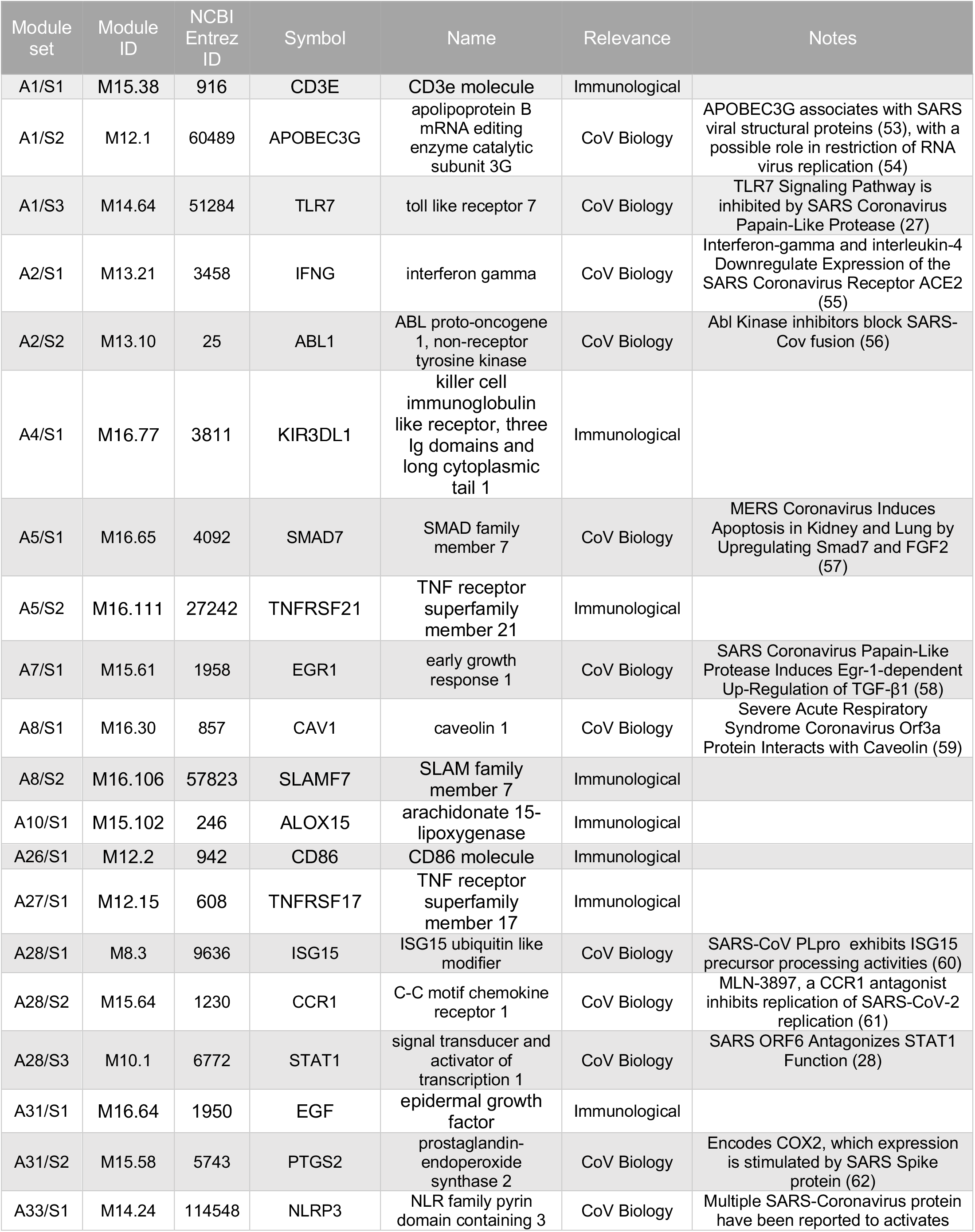
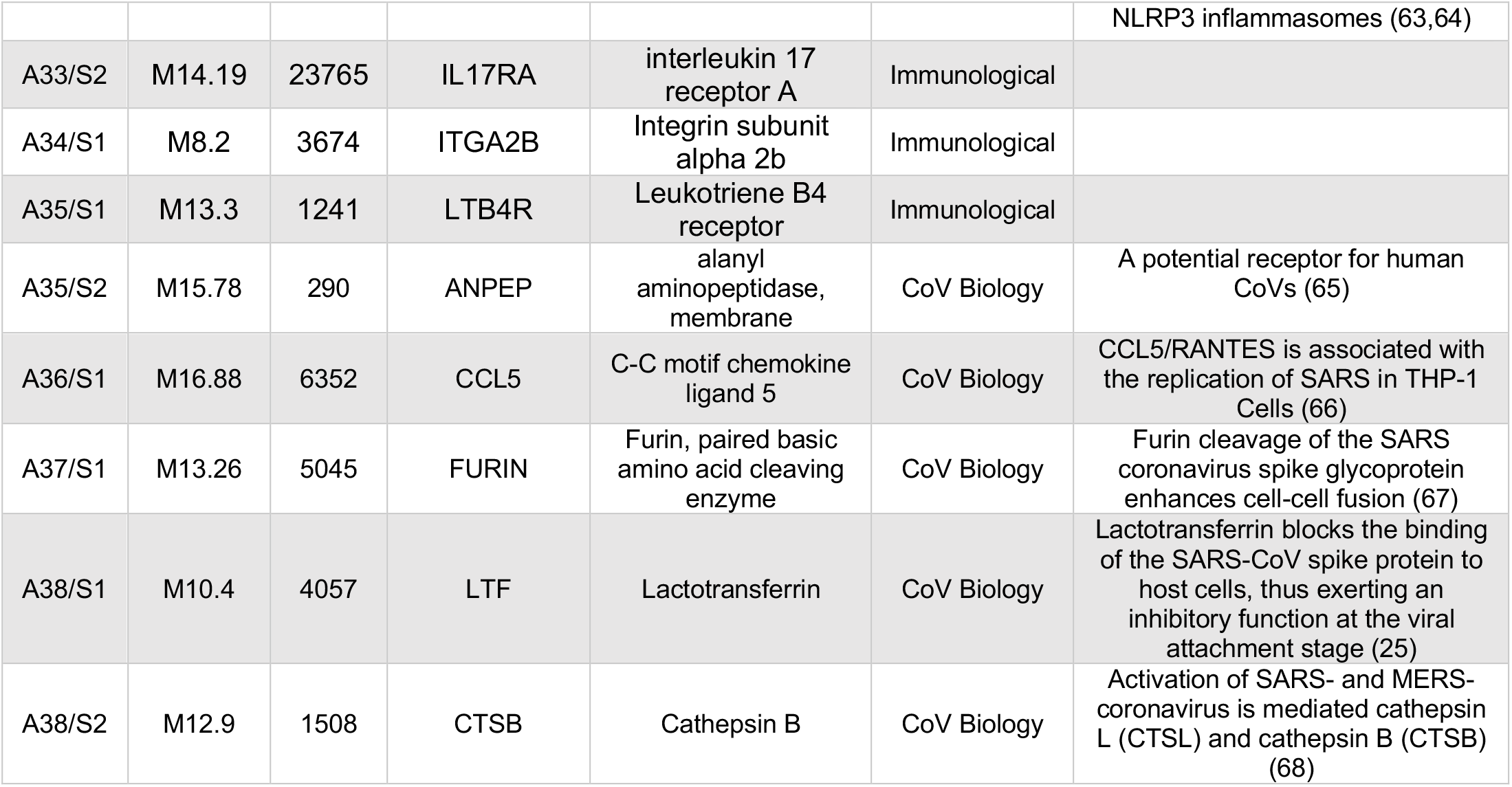
Illustrative targeted panel – SARS biology relevance focus

This screen identified several molecules that may be of importance for SARS-CoV-2 entry and replication. It is expected that this knowledge will evolve rapidly over time and frequent updates may be necessary. And, as for the previous two panels, investigators may also have an interest in including more than one candidate per module set. This of course would also be feasible, although at the expense of course of parsimony.

### Development of an annotation framework in support of signatures curation efforts

A vast amount of information is available to support the work of expert curators who are responsible for finalizing the selection of candidates. This process often requires accessing a number of different resources (e.g. those listed in **Table 2**). Here we have built upon earlier efforts to aggregate this information in a manner that makes it seamlessly accessible by the curators.

As proof of principle, we created dedicated, interactive presentations in Prezi for module aggregates A28 ((24)) and A31 (https://prezi.com/view/zYCSLyo0nvJTwjfJkJqb/). These presentations are intended, on the one hand, to aggregate contextual information that can serve as a basis for data interpretation. On the other hand, they are intended to capture the results of the interpretative efforts of expert curators.

The interactive presentations are organized in sections, each showing aggregated information from a different level: module-sets, modules and transcripts (**Figure 5**). The information derived from multiple online sources, including both third party applications and custom applications developed by our team (**Table 2**). Among those is a web application developed specifically for this work, which was used to generate the Covid-19 plots from Ong *et al.* and Xiong *et al.* (**Figure 5A**). The interactive presentation itself permits to zoom in and out, determine spatial relationships and interactively browse the very large compendium of analysis reports and heatmaps generated as part of these annotation efforts. The last section that contains transcript-centric information, is also the area where interpretations from individual curators is aggregated.

**Figure 4.**
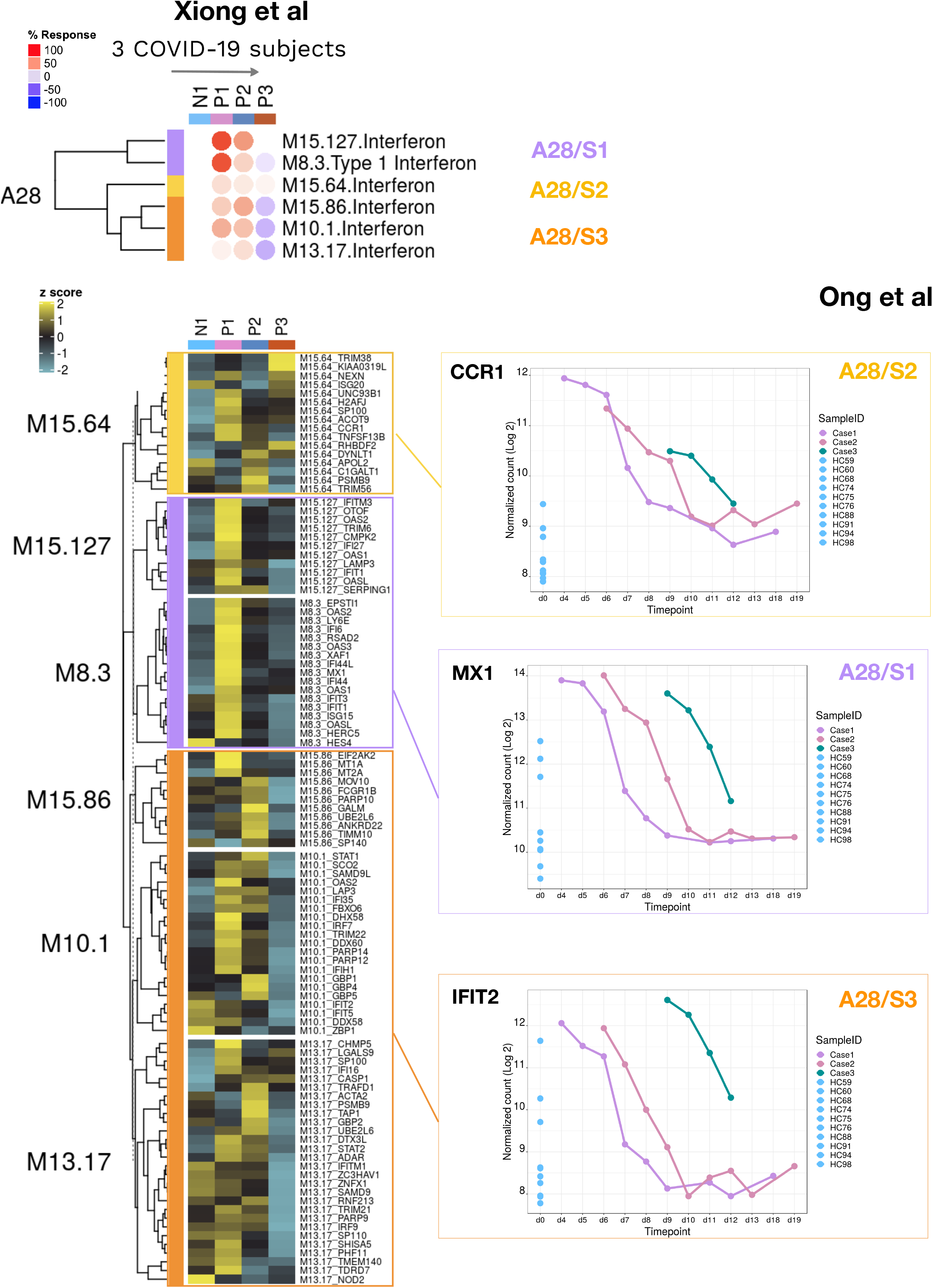
Changes in abundance of transcripts comprising aggregate A28 in response to SARS-CoV-2 infection. The heatmaps display the changes in transcript abundance in three Covid-19 patients comprising the Xiong *et al.* RNA-seq transcriptome datasets. The top heatmap summarizes the module-level values for the six modules forming aggregate A28. The color code indicates membership to one of the three Covid-19 module sets that were defined earlier. The bottom heatmap shows patterns of abundance for the same six modules, but at the individual gene level. The line graphs on the right show changes in abundance for three transcripts from the “therapeutic relevance panel” in three Covid-19 patients profiled by Ong *et al*. using a generic Nanostring immune set comprising 594 transcripts.

**Figure 5:**
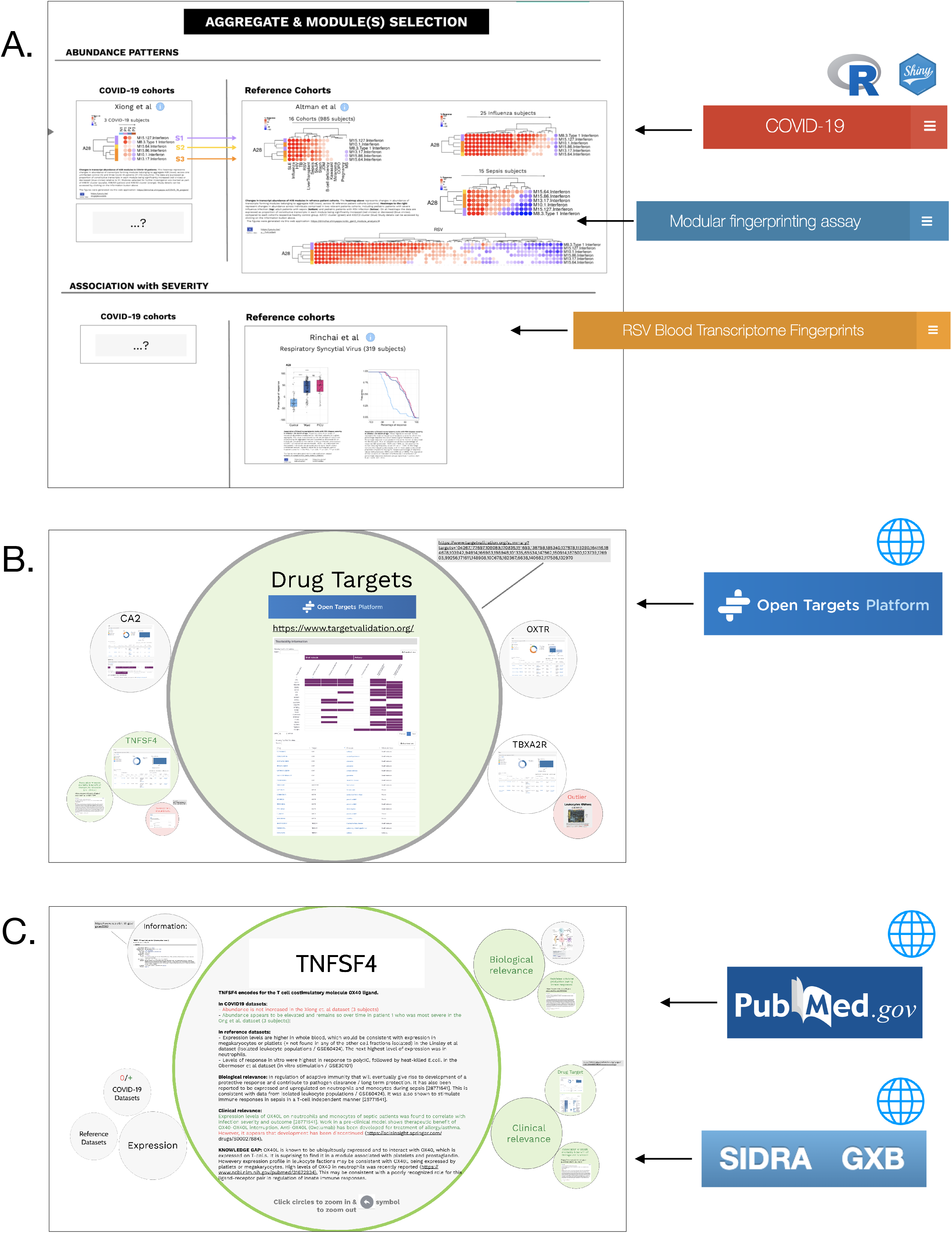
High resolution annotation framework supporting the curation and interpretation of Covid-19 module sets. This series of screenshots shows the content of the interactive presentations that have been established to provide curators with access to detailed annotations regarding modules forming a given aggregate, its constitutive modules and targets that have been selected for inclusion in transcript panels. Links to interactive presentations and resources mentioned below are available in **Table 2**. **A. Module aggregate-level information.** This section displays patterns of transcript abundance across the modules forming a given aggregate, as well as the degree of association of this aggregate with the severity of RSV disease. Plots used to populate this section were generated using three web applications, including one that was developed in support of this work that compiles the Covid-19 blood transcriptional data available to date. The other two applications were developed as part of a previous study to generate plots for the reference disease cohorts and RSV severity association plots (45). **B. Module-level information.** This section includes, for a given module, reports from functional profiling tools as well as patterns of transcript abundance across the genes forming the module. Drug targeting profiles were added to provide another level of information. **C. Gene-centric information.** The information includes curated pathways from the literature, articles and reports from public resources. Gene-centric transcriptional profiles that are available via gene expression browsing applications deployed by our group are also captured and used for context (GXB). A synthesis of the information gathered by expert curation and potential relevance to SARS-Cov-2 infection can also be captured and presented here.

We have annotated and interpreted some of the transcripts included in **A31/S1** in such a manner: **1) OXTR,** which encodes for the Oxytocin receptor through which antiinflammatory and wound healing properties of Oxytocin are mediated (29). Among our reference cohort datasets, OXTR is most highly increased in patients with *S. aureus* infection or active pulmonary tuberculosis (7). **2) CD9,** which encodes a member of the tetraspanin family, facilitates the condensation of receptors and proteases activating MERS-CoV and promoting its rapid and efficient entry into host cells (30). **3) TNFSF4**, which encodes for OX40L and is a member of the TNF superfamily. Although OX40L is best known as a T-cell co-stimulatory molecule, reports have also shown that it is present on the neutrophil surface (31). Furthermore, OX40L blockade improved outcomes of sepsis in an animal model.

Our interpretation efforts have been limited thus far by expediency. Certainly, interpretation will be the object of future, more targeted efforts. In the meantime, this annotation framework supports the selection of candidates forming the panels presented here. It may also serve as a resource for investigators who wish to design custom panels of their own.

## DISCUSSION

Early reports point to profound immunological changes occurring in affected patients during the course of a SARS-CoV-2 infection (32,33). In particular, patterns of immune dysfunction have been associated with clinical deterioration and the onset of severe respiratory failure (34). However, disease outcomes remain highly heterogeneous and factors contributing to clinical deterioration are poorly understood. Among other modalities, means to establish comprehensive immune monitoring in cohorts of Covid-19 patients are needed.

Here we designed an approach select and curate targeted blood transcript panels relevant to Covid-19. When finalized, such panels could in turn serve as a basis for rapid implementation of focused transcript profiling assays. This process should become possible as more Covid-19 blood transcriptome profiling datasets become available in the coming weeks and months. These data could, for instance, be used to refine the delineation of Covid-19 module sets.

Because our selection strategy relies primarily on a pre-existing module repertoire framework, we anticipate that changes would only be relatively minor. Indeed, one advantage of basing candidate selection on a repertoire of transcriptional modules is that it permits to derive non-synonymous transcripts sets. In other words, each transcript included in the panel could survey the abundance of a different module (signature). Basing selection on differential expression instead, for instance, would tend to select multiple transcripts from more dominant signatures (with highest significance / fold changes). But for more specific purposes, such as differential diagnosis or prediction, machine learning models would be more appropriate [for instance in sepsis studies: (35)]. Indeed, such panels have already been developed for Covid-19 (36), and we anticipate that more will emerge over the coming months. However, our intent here was different: our primary aim was to support the development of a solution that can monitor immune responses and functions.

Delineation of “immune trajectories” associated with clinical worsening of Covid-19 patients is one application to consider. Another application would be the measurement of responses to therapy (as part of standard of care or a trial). The immune profiling of asymptomatic or pre-symptomatic patients (e.g quarantined) would be another setting where implementation of such an assay could prove useful. For this, it would for instance be possible to use protocols that we have previously developed for home-based, self-sampling and blood RNA stabilization (37,38).

Different connotations were given for the three panels, which are presented here as a proof of principle. The panel consisting of immunologically relevant markers might have the highest general interest. However, measuring changes in the abundance of transcripts coding for molecules that are targetable by existing drugs could have higher translational potential. Another illustrative panel comprises transcripts coding for molecules that are of relevance to SARS-CoV-2 biology and might be of additional interest in investigations of host–pathogens interactions. The common denominator between these panels is that they comprise representative transcripts of each of the 28 module sets. Other transcript combinations following the same principle would be possible, as well as the inclusion of multiple transcripts from the same set for added robustness. The obvious disadvantage of the latter, however, is the increase to the size of the panel. Medium-throughput technology platforms, such as the Nanostring Ncounter System, Fluidigm Biomark or ThermoFisher Openarray, would be appropriate for implementing custom profiling assays with the number of targets comprising the tentative panels presented here (or a combination thereof). Downsizing panels to comprise ±10 key markers might serve as a basis for implementation on more ubiquitous real-time PCR platforms.

Overall, this work lays the ground for a framework that could support the development of increasingly more refined and interpretable targeted panels for profiling immune responses to SARS-CoV-2 infection. This should be possible in part through the further development of environments providing investigators with seamless access to vast amounts of annotations aggregated from different sources.

## ACKNOWLEDGEMENTS

Development of some of the bioinformatic resources and approaches employed here was supported by NPRP grant # 10-0205-170348 from the Qatar National Research Fund (a member of Qatar Foundation). The work reported herein is solely the responsibility of the authors.

## AUTHOR CONTRIBUTIONS

DR, BK, MT, NB, Malt, MB, GZ, ADM, BT, DB, DC conceptualization. DR, BK, MT, ZC, SD, TB, MG: data curation and validation. DR, BK, MT and DC: visualization. DR, BK, MT, ZC, SD, TB, MG, DB, DC: analysis and interpretation. DB, DC: funding acquisition. DR and DC: methodology development. DR, BK, MT, DC: writing of the first draft. DR, MT, MAlf: software development and database maintenance. DR, BK, MT, ZC, SD, TB, MG, RB, NB, MAlf, Malt, AB, MB, GZ, ADM, BT, DB, DC writing–review and editing The contributor’s roles listed above follow the Contributor Roles Taxonomy (CRediT) managed by The Consortia Advancing Standards in Research Administration Information (CASRAI) (https://casrai.org/credit/).

---

**Supplementary Figure 1:**
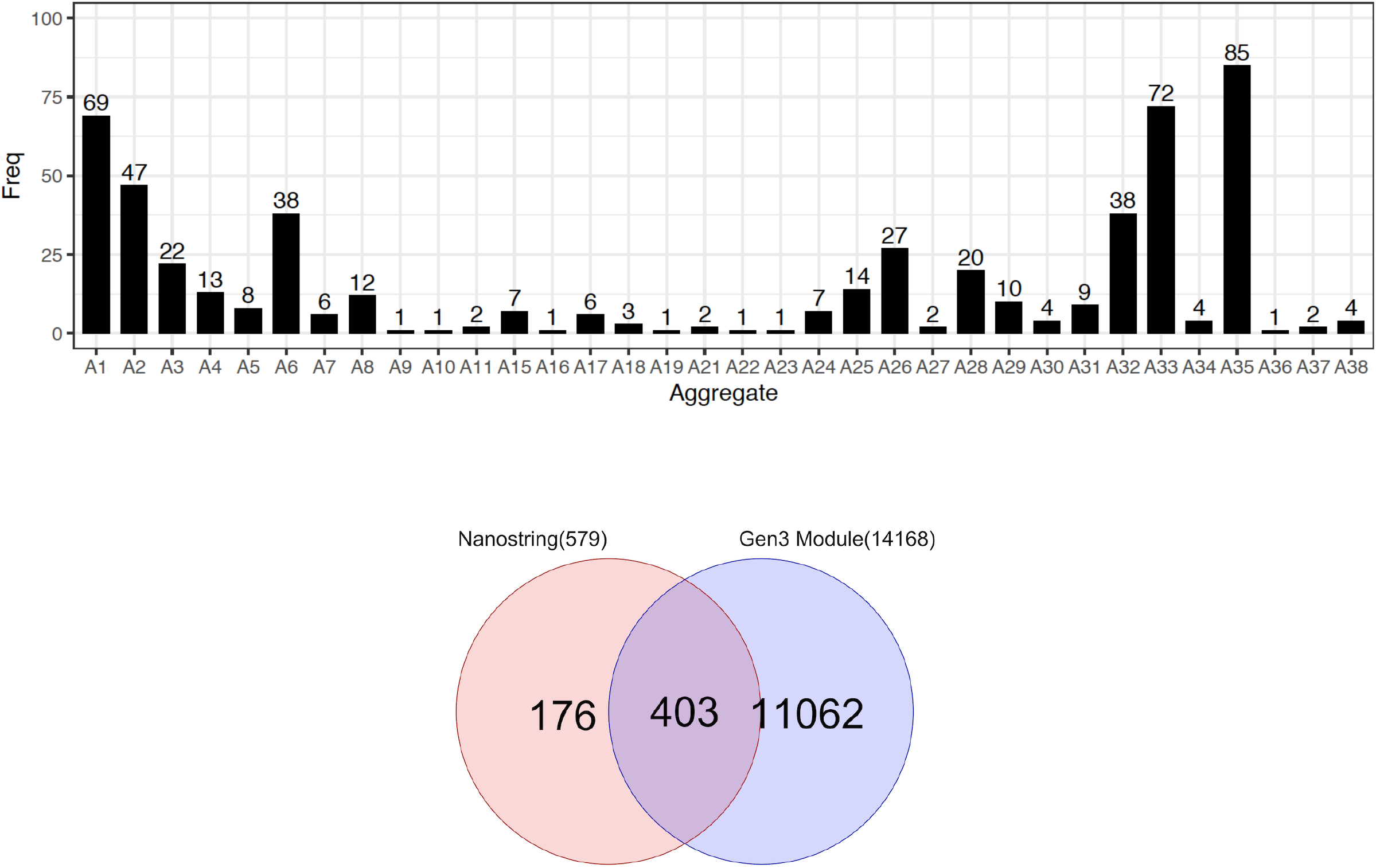
Coverage of the pre-established 38 transcriptional module aggregate repertoire by the Nanostring immunology panel 2. The bar graphs show the distribution of the 579 transcript constituting the standard Nanostring immunology panel used by Ong *et al.* across the 38 module aggregates forming this repertoire. The Venn diagram shows the degree of overlap between the Nanostring panel and the transcripts forming this modular repertoire.

## METHODS

### Datasets

Two Covid-19 blood transcriptional datasets available at the time this work was conducted were used: **1)** Xiong *et al.* (9) obtained peripheral mononuclear cell samples obtained from one uninfected control individual and three patients with Covid-19. RNA abundance was profiled via RNAseq. The data were deposited in the Genome Sequence Archive of the Beijing Institute of Genomics, Chinese Academy of Sciences, under the accession number CRA002390. FASTQ passed QC and were aligned to reference genome GRChg38/hg19 using Hisat2 (v2.05). BAM files were converted to a raw count expression matrix using subreads (v1.6.2). Raw expression data was corrected for within lane and between lane effects using R package EDASeq (v2.12.0) and quantile normalized using preprocessCore (v1.36.0). The modular analysis was performed by using 10,617 RNA-seq genes which overlapped with transcripts from the 3rd generation module construction (7). details of the analysis as described below section.

**2)** Ong *et al.* (8) collected whole blood stabilized in RNA buffer from uninfected controls and three Covid-19 patients at multiple time points. RNA abundance was profiled using a standard immunology panel from Nanostring comprising 594 transcripts. The data were deposited in the arrayexpress public repository with accession ID E-MTAB-8871. The normalized data were downloaded, and modular analysis was performed by using 403 NanoString genes which overlapped with transcripts from the 3nd generation module construction details of the analysis as described below section.

**3)** We also used a reference dataset generated by our group that was previously used to construct the 382 blood transcriptional module repertoire (7). Briefly, this repertoire consists of the following cohorts of patients and respective control subjects: *S. aureus* infection (99 cases, 44 controls), sepsis (35 cases, 12 controls), tuberculosis (23 cases, 11 controls), Influenza (25 cases, 14 controls), RSV infection (70 cases, 14 controls), HIV infection (28 cases, 35 controls), systemic lupus erythematosus (55 cases, 14 controls), multiple sclerosis (34 cases, 22 controls), juvenile dermatomyositis (40 cases, 9 controls), Kawasaki disease (21 cases, 23 controls), systemic onset idiopathic arthritis (62 cases, 23 controls), COPD (19 cases, 24 controls), melanoma (22 cases, 5 controls), pregnancy (25 cases, 20 controls), liver transplant recipients (94 cases, 30 controls), and B-cell deficiency (20 cases, 13 controls). All samples were run at the same facility on Illumina HumanHT-12 v3.0 Gene Expression BeadChips. The data have been deposited in NCBI GEO with accession number GSE100150.

### Transcriptional module repertoire

The method used to construct the transcriptional module repertoire has been described elsewhere (39,40). The version used here is the third and last to have been developed by our group over a period of 12 years. It is the object of a separate publication (available on a pre-print server (7)).

Briefly, the approach consists of identifying sets of co-expressed transcripts in a wide range of pathological or physiological states, focusing in this case on the blood transcriptome as the biological system. We determined co-expression based on patterns of co-clustering observed for all gene pairs across the collection of 16 reference datasets listed in the previous section and that encompassed viral and bacterial infectious diseases as well as several inflammatory or autoimmune diseases, B-cell deficiency, liver transplantation, stage IV melanoma and pregnancy. Overall, this collection comprised 985 blood transcriptome profiles. A weighted, co-expression network was built with the weight of the nodes connecting a gene pair being based on the number of times co-clustering was observed for the pair among the 16 reference datasets. Thus, the weights ranged from 1 (where co-clustering occurs in one of 16 datasets) to 16 (where co-clustering occurs in all 16 datasets). Next, this network was mined using a graph theory algorithm to define subsets of densely connected gene sets that constituted our module repertoire (“Cliques” and “Paracliques”).

Overall, 382 transcriptional modules were identified, encompassing 14,168 transcripts. A supplemental file including the definition of this module repertoire along with the functional annotations is available elsewhere (7). To provide another level of granularity and facilitate data interpretation, a second round of clustering was performed to group the modules into “aggregates”. This process was achieved by grouping the set of 382 modules according to the patterns of transcript abundance across the 16 reference datasets that were used for module construction. This segregation resulted in the formation of 38 aggregates, each comprising between one and 42 modules.

### Module repertoire analyses

The modular analyses were performed using the core set of 14,168 transcripts forming the module repertoire. For group-level comparisons (cases vs controls), a paired t-test was performed on the log2-transformed data [Fold change (FC) cut off = 1.5; FDR cut off = 0.1]. For individual-level analyses, each sample was compared to the mean value of the corresponding control samples (or individual sample in the case of the Xiong *et al.* dataset). The cut off comprised an absolute FC >1.5 and a difference in counts >10. The results for each module are reported as the percentage of its constitutive transcripts that increased or decreased in abundance. Because the genes comprised in a module are selected based on the co-expression observed in blood, the changes in abundance within a given module tend to be coordinated and the dominant trend is therefore selected (the greater value of the percentage increased vs. percentage decreased). Thus, the values range from −100% (all constitutive modules are decreased) to +100% (all constitutive modules are increased). A module was considered to be “responsive” when the proportion of transcripts found to be increased was >15%, or when the proportion of transcripts found to be decreased was ≤15%. At the aggregate-level, the percent values of the constitutive modules were averaged.

### Data visualization

Changes in transcript abundance reduced at the module or module aggregate-level were visualized using a custom fingerprint heatmap format. For each module, the percentage of increased transcripts is represented by a red spot and the percentage of decreased transcripts is represented by a blue spot. The fingerprint grid plots were generated using “ComplexHeatmap” (41). A web application was developed to generate the plots and browse modules and module aggregates (https://drinchai.shinyapps.io/COVID_19_project/). A detailed description and source code will be available as part of a separate publication BioRxiv deposition on GitHub and BioRxiv (in preparation).

### Selection of transcripts for inclusion in targeted panels

#### Therapeutic relevance

Covid-19 module sets belonging to aggregates comprising module annotations relating to inflammation, monocytes, neutrophils or coagulation pathway were selected for screening (A7, A8, A26, A31, A33, A34, A35). In turn, transcripts from each of the corresponding module sets were selected on the basis of their status as a known therapeutic target of a drug for which clinical precedence exists (source: targetvalidation.org). Next, candidates were prioritized via expert curation on the basis of compatibility and a potential benefit as a Covid-19 treatment. Curators were tasked with prioritizing candidates within each of the Covid-19 module sets. Only the top ranked candidate from each set was selected for inclusion in the panel. Module sets from aggregate A28 (interferon response) may also be of clinical relevance, may also be of clinical relevance, as indicators of a treatment response since interferon administration has been shown to increase the activity of anti-viral drugs in Covid-19 patients (42). The selection of candidates for aggregate A28 sets was thus based on the amplitude of the response to beta-interferon therapy measured in patients with multiple sclerosis [fold-change over pretreatment baseline (43) & NCBI GEO accession GSE26104]. The remaining nine aggregates, which tended to associate preferentially with adaptive immune responses and for which targeting by therapies might prove detrimental, were not included in this screen. For these, representative transcripts from the default panel of immune relevant transcripts were included.

#### Relevance to Coronavirus biology

for the second panel, transcripts were primarily selected based on their relevance to SARS biology. As a first step, a literature profiling tool was used to identify among the SARS, MERS, or Covid-19 literature articles that were associated with transcripts forming the 28 Covid-19 module sets (Literature Lab, Gene Profiler module; Accumenta Biotech, Boston, MA). Next, the potential associations were assessed by manual curation. The curators prioritized the transcripts for which the associations could be confirmed based on importance and robustness.

#### Immunological relevance

Lists of immunologically relevant genes were retrieved from Immport (44), and were used along with membership to IPA pathways (Ingenuity Pathway Analysis, QIAGEN, Germantown MD) to annotate transcripts comprising Covid-19 module sets. The curators prioritized annotated transcripts on the basis of their relevance to the functional annotations of the module set (e.g. interferon, inflammation, cytotoxic cells). The transcript with the highest priority rank was included in the assay.

#### Housekeeping genes

A recommended set of housekeeping genes is provided in **Table 6**. These were selected on the basis of low variance observed across the 985 transcriptome profiles generated for our reference cohorts.

**Table 6:**
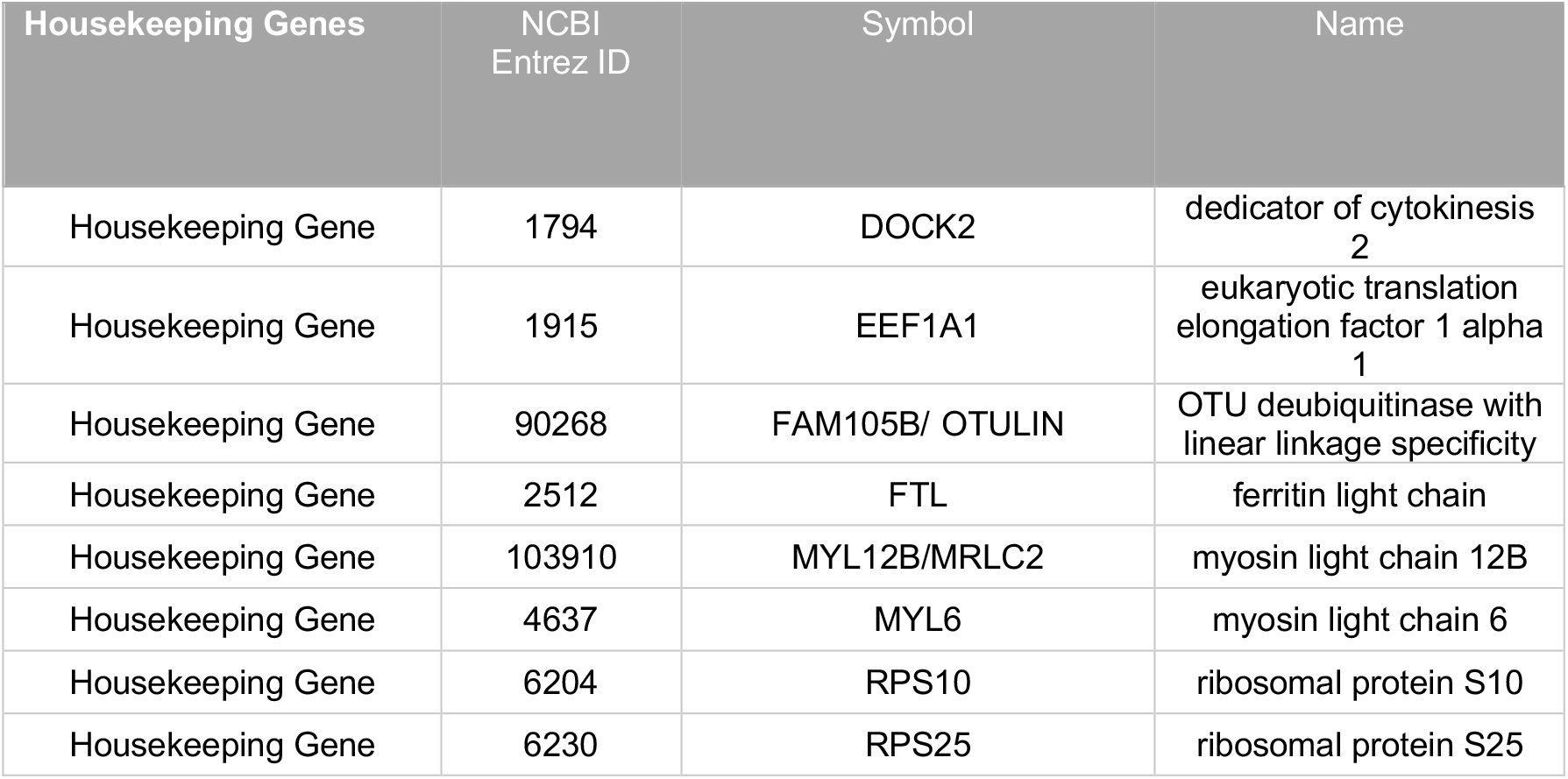
List of housekeeping genes that may be suitable for blood transcript profiling applications

### Annotation framework

Links to the resources described in this section and to video demonstrations are available in **Table 2.** Interactive presentations were created via the Prezi web application. For this we have built and expanded upon an annotation framework established as part of the characterization of our reference blood transcriptome repertoire (7). Several bioinformatic resources were used to populate interactive presentations that served as a framework for annotation of Covid-19 relevant module sets. These resources include web applications deployed using Shiny R, which permit to plot transcript abundance patterns at the module and aggregate levels. Two of these applications were developed as part of a previous work establishing the blood transcriptome repertoire and applying it in the context of a metaanalysis of six public RSV datasets (12). A third application was developed as part of this work and can generate profiles at the transcript, module and module-aggregate levels for the Xiong *et al.* and Ong *et al.* datasets.

## SUPPLEMENTAL INFORMATION

Supplemental File 1: Delineation of Covid-19 relevant modules sets in all 17 aggregates retained in the first step of the selection process.

