## Supplementary File 1 for "A modular framework for the development of targeted Covid-19 blood transcript profiling panels"

**Delineation of Covid-19 relevant modules sets. The** heatmaps on the left presented in this file represents the abundance levels for transcripts forming modules belonging to a given aggregate (rows), across three Covid-19 patients (P1-P3) relative to one uninfected control subject (columns) [Xiong et al dataset]. The data are expressed as the proportion of constitutive transcripts in each module being significantly increased (red circles) or decreased (blue circles) relative to N1. The heatmaps on the right represent the abundance levels for transcripts forming the same set of modules across 16 reference patient cohorts (columns).

### Aggregate A1: Covid-19 relevant sets S1, S2 & S3

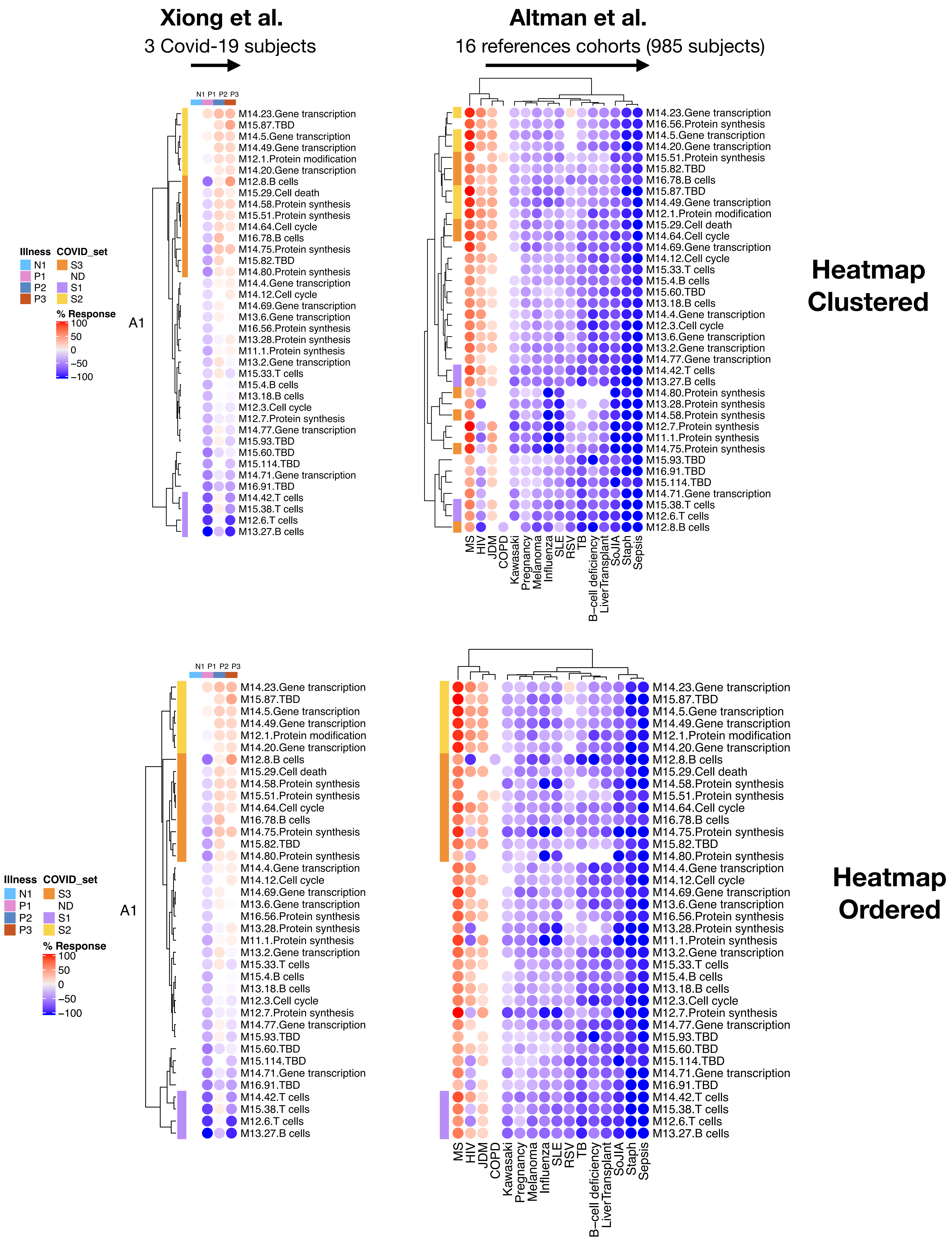

### Aggregate A2: Covid-19 relevant sets S1 & S2

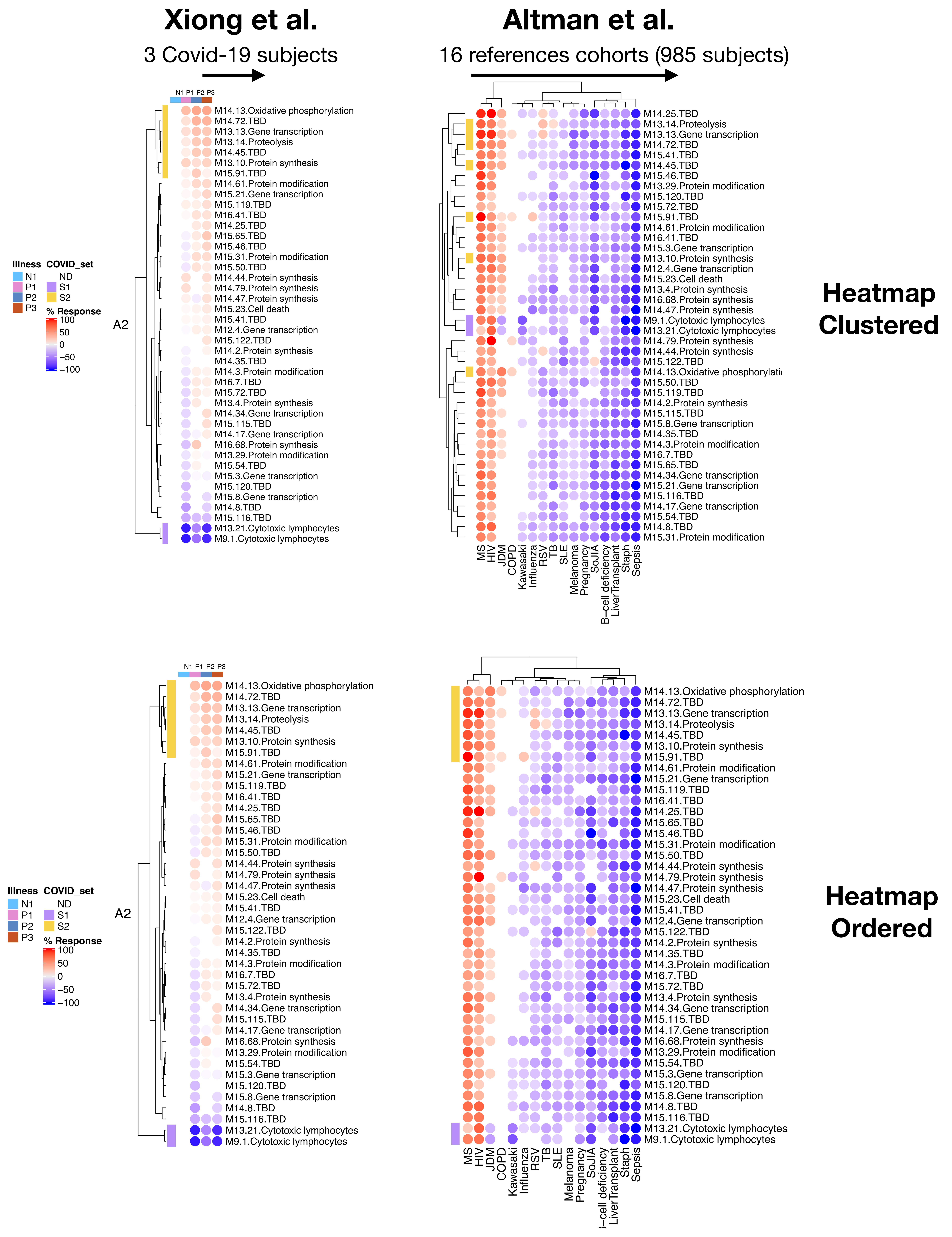

### Aggregate A4: Covid-19 relevant sets S1

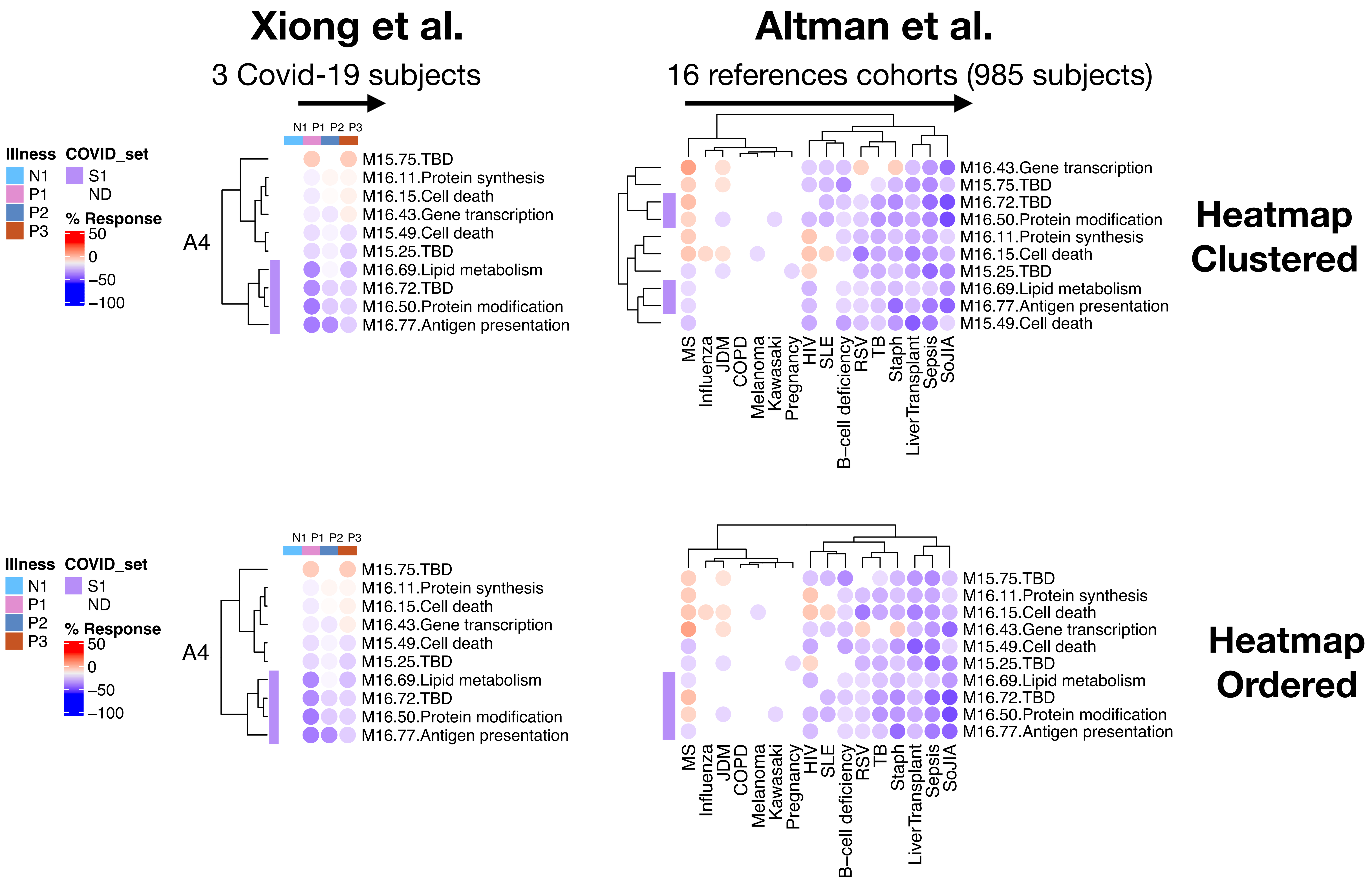

### Aggregate A5: Covid-19 relevant sets S1 & S2

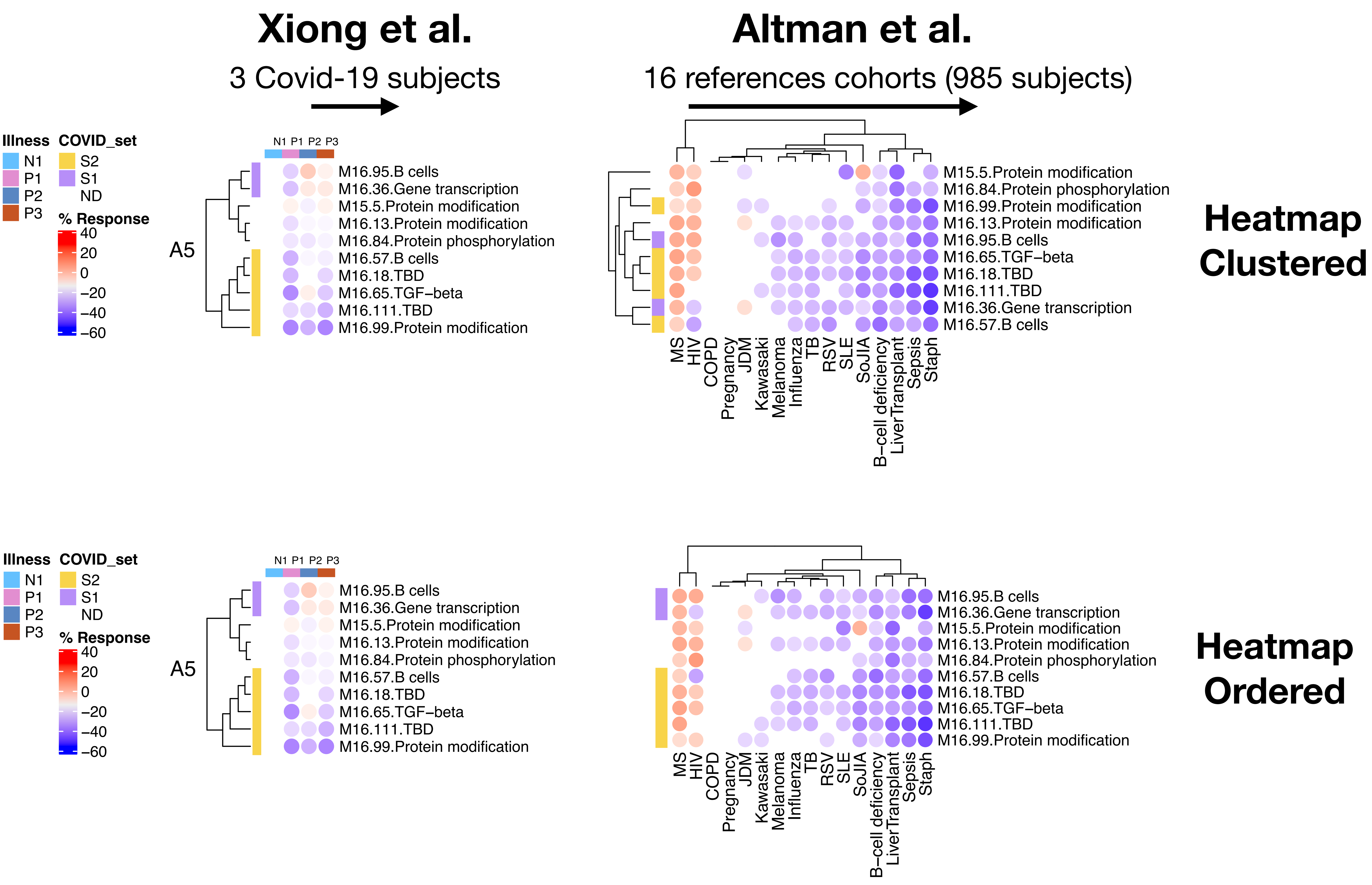

### Aggregate A7: Covid-19 relevant sets S1

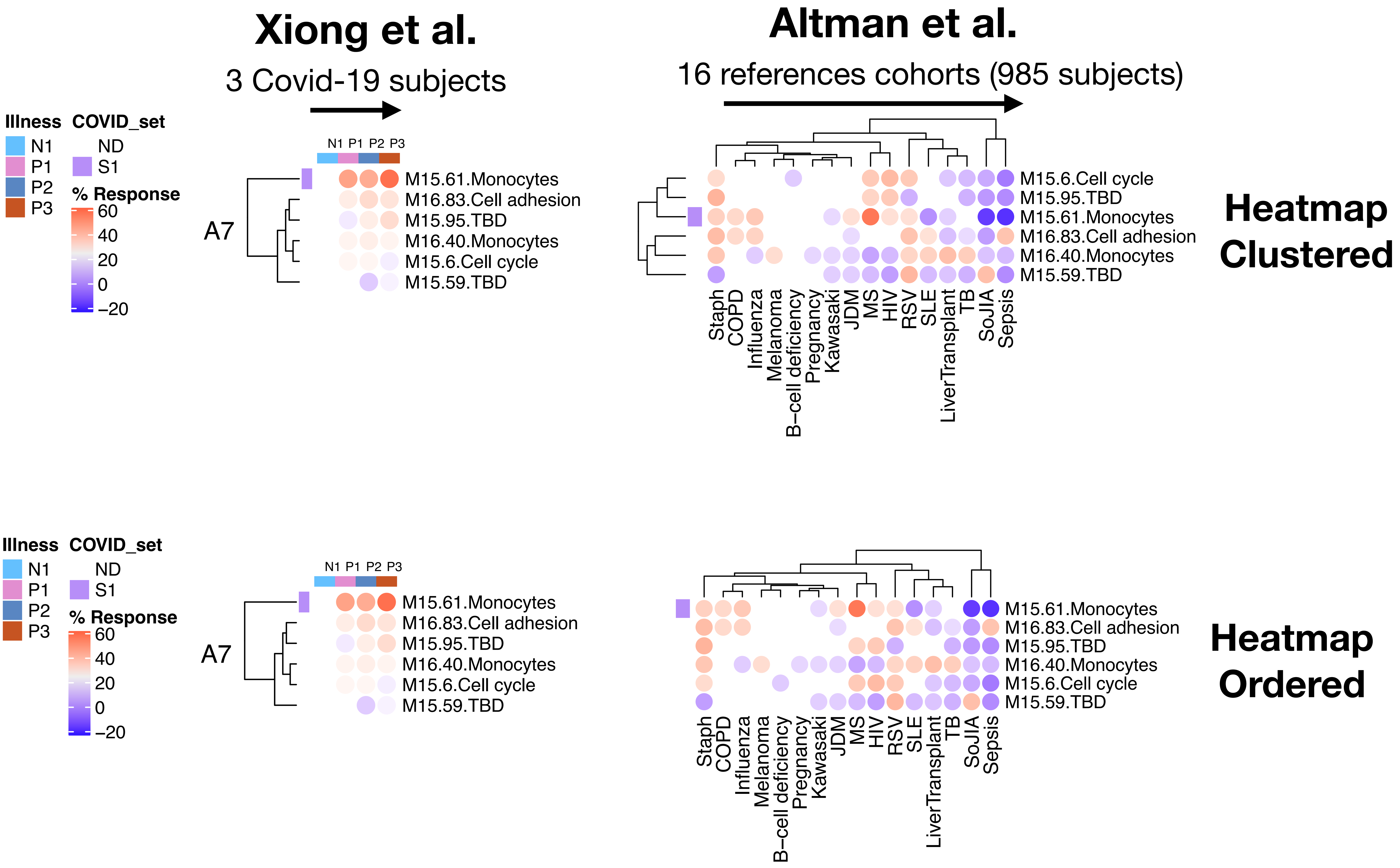

### Aggregate A8: Covid-19 relevant sets S1 & S2

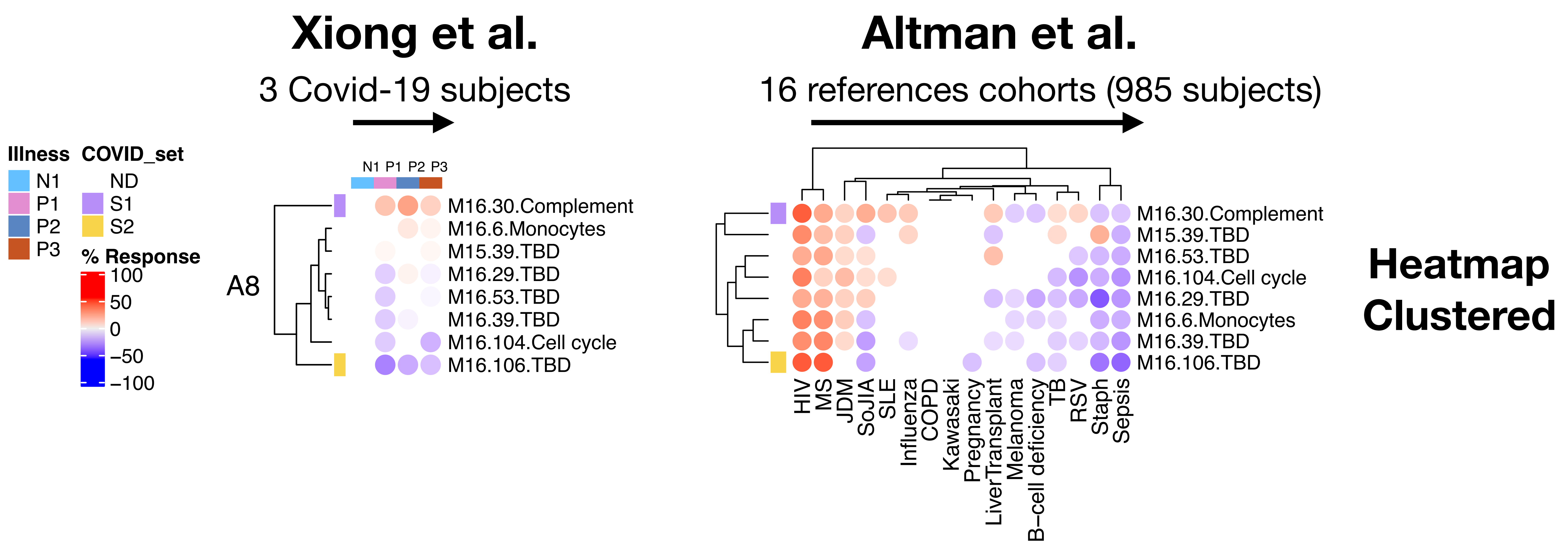


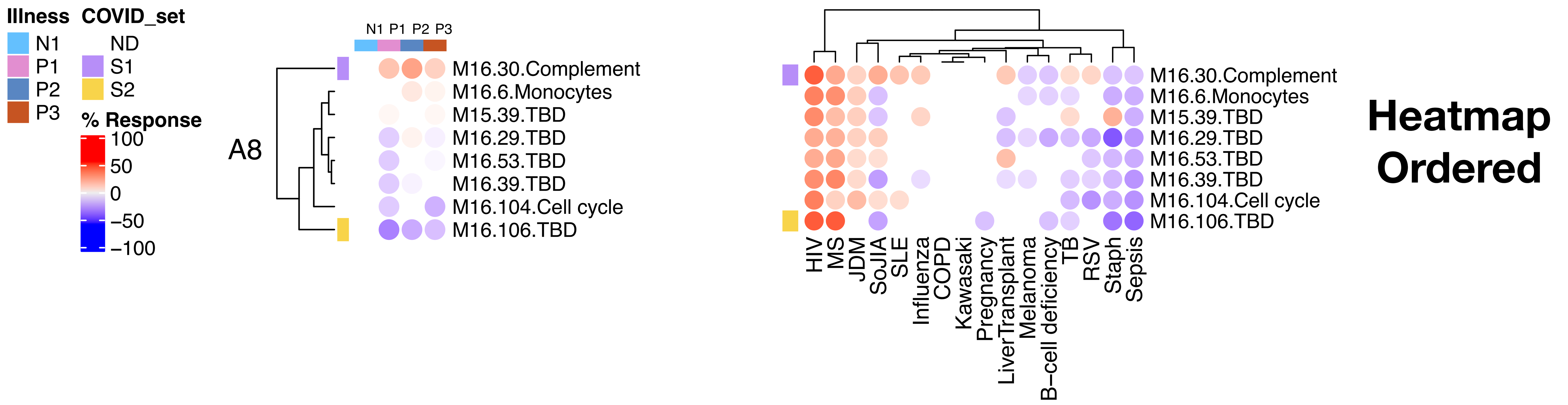

### Aggregate A10: Covid-19 relevant sets S1

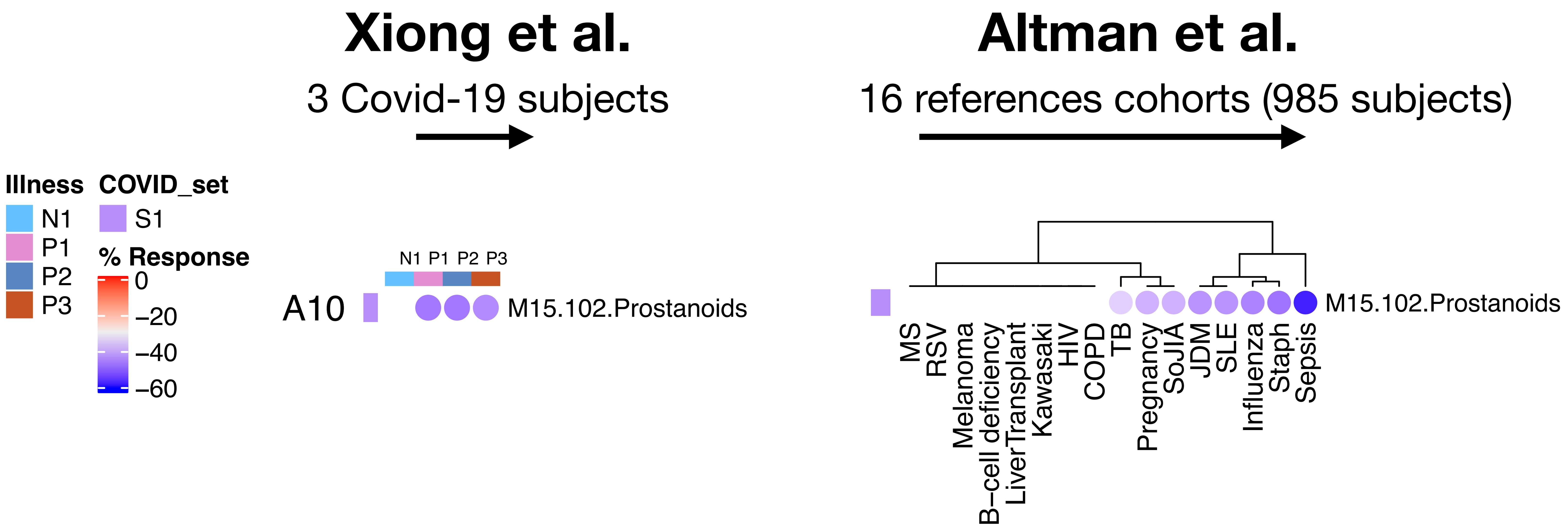

### Aggregate A26: Covid-19 relevant sets S1

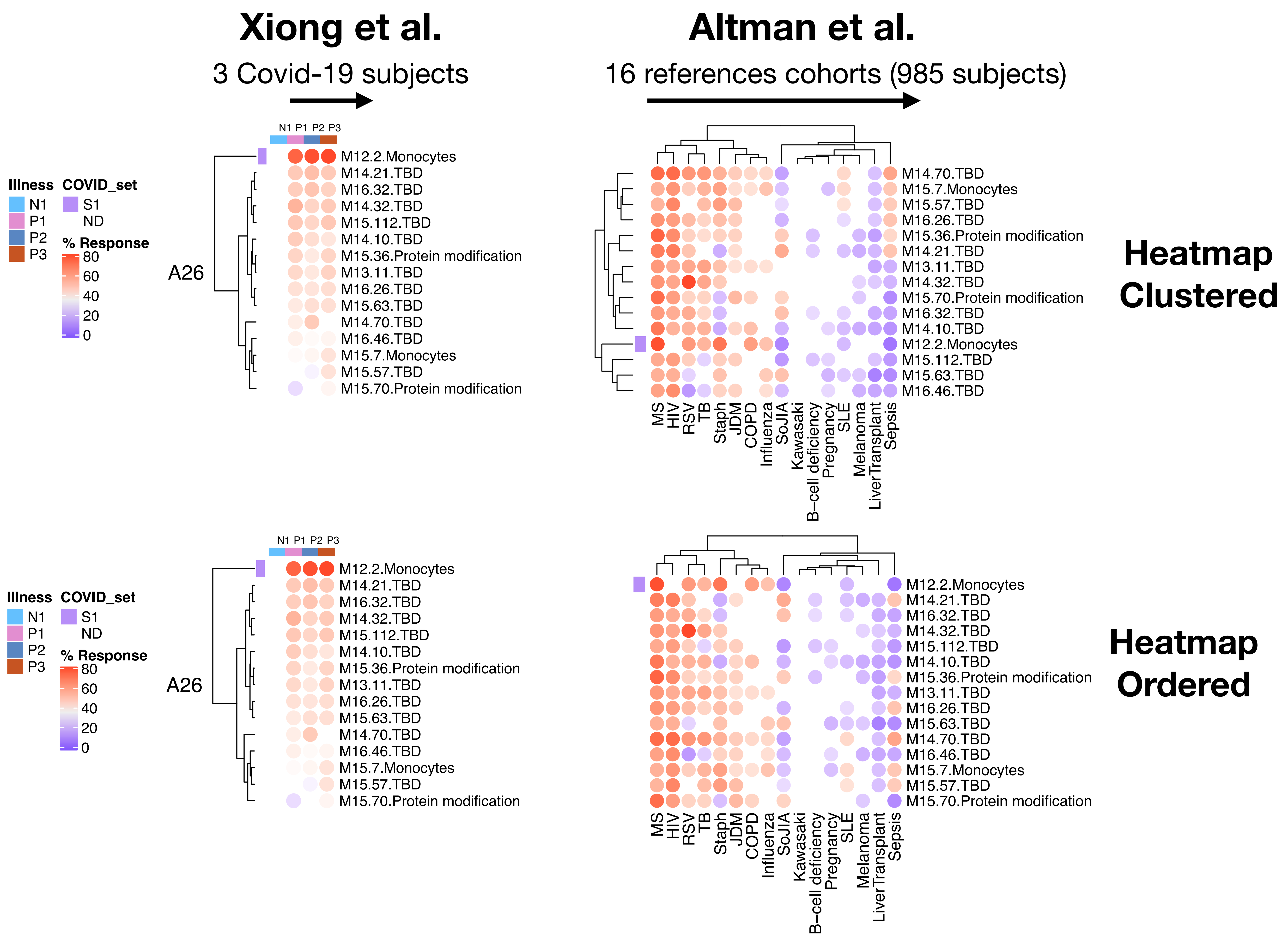

### Aggregate A27: Covid-19 relevant sets S1

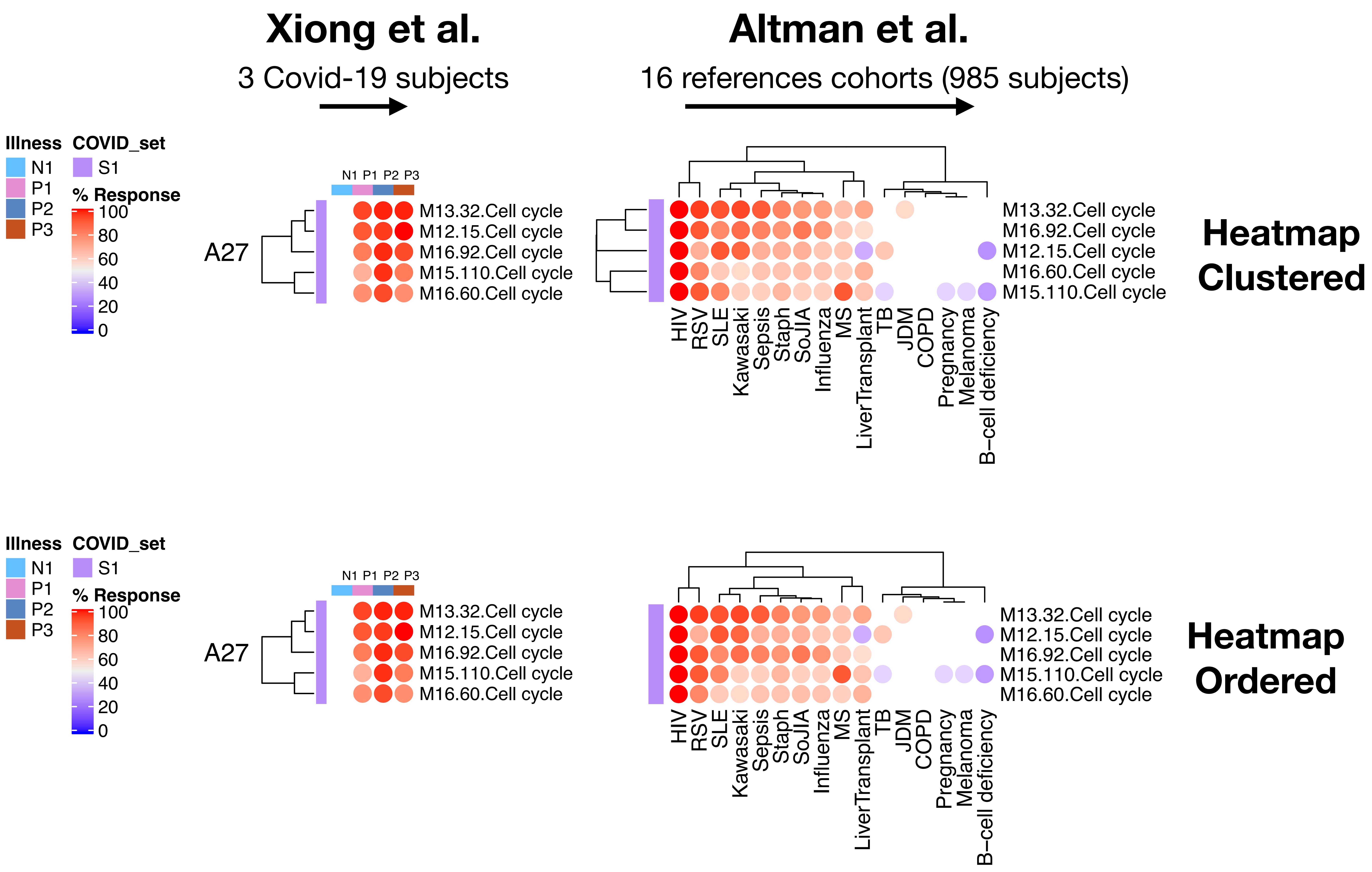

### Aggregate A28: Covid-19 relevant sets S1, S2 & S3

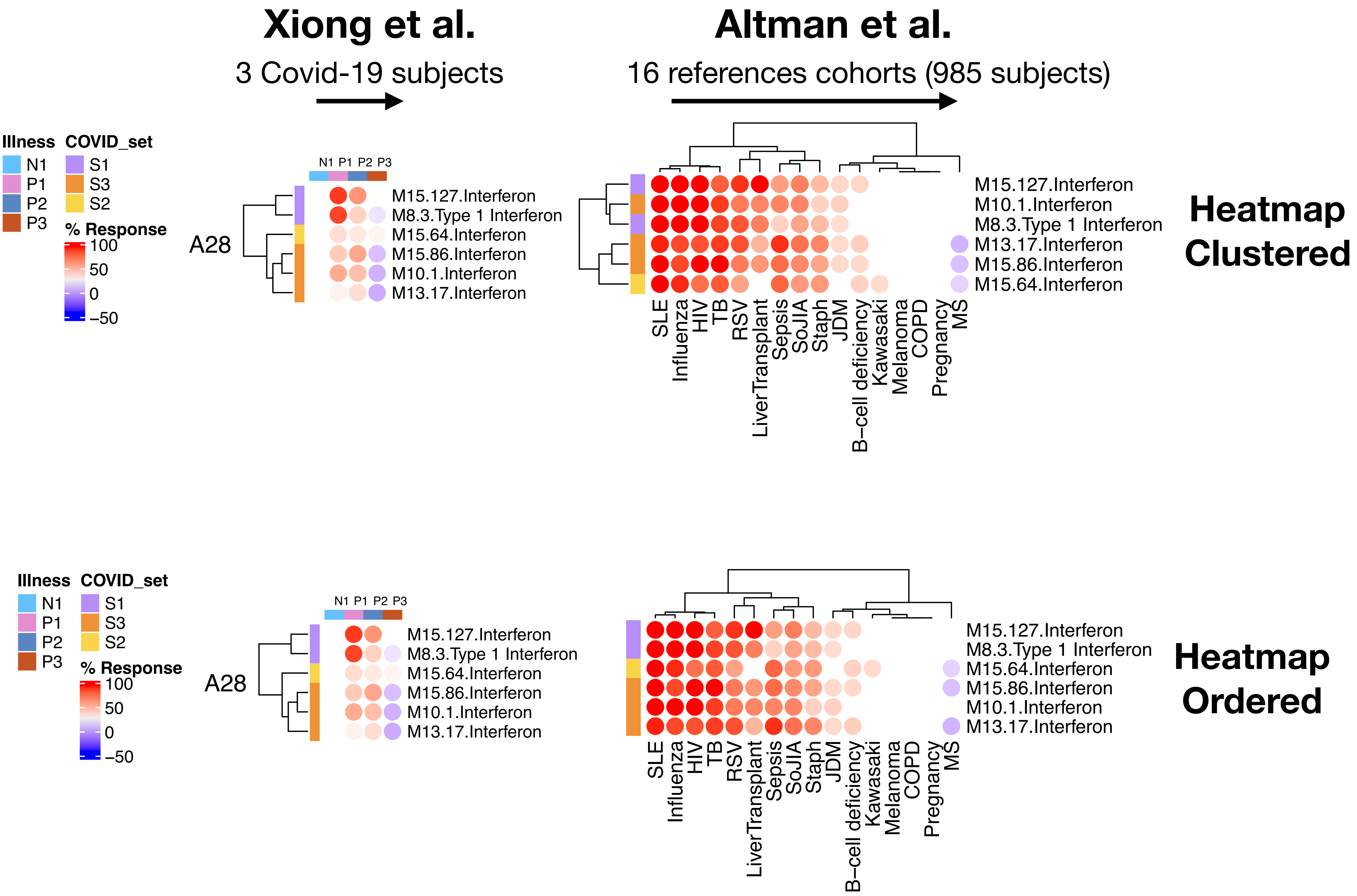

### Aggregate A31: Covid-19 relevant sets S1 & S2

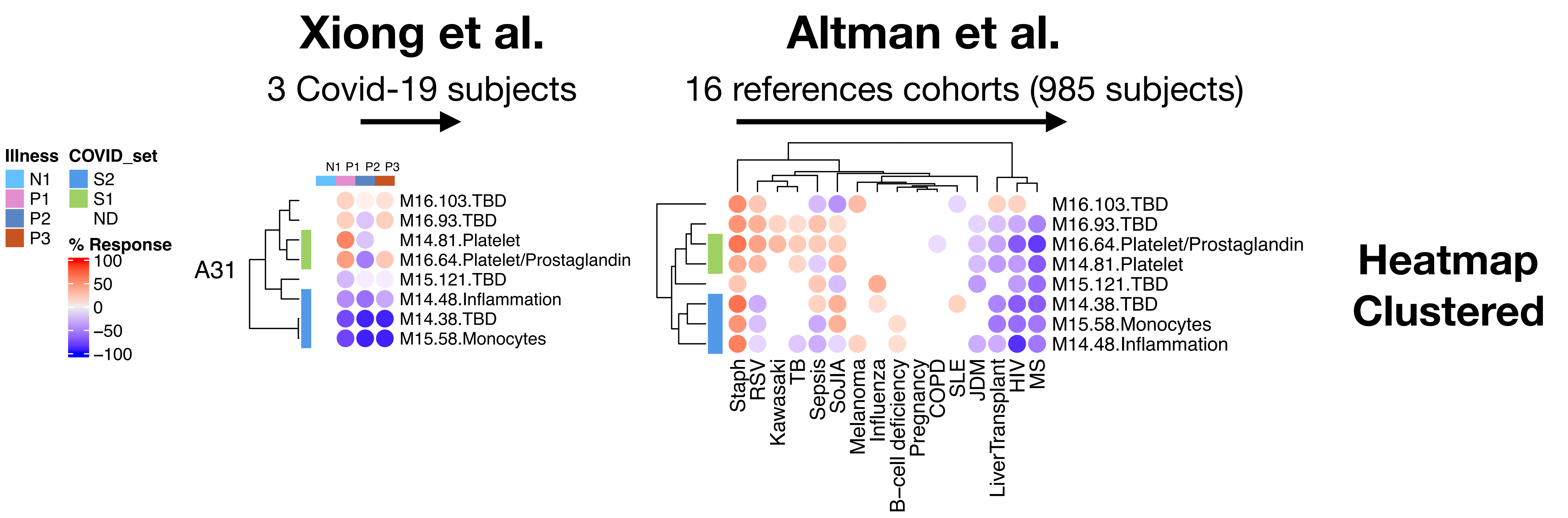


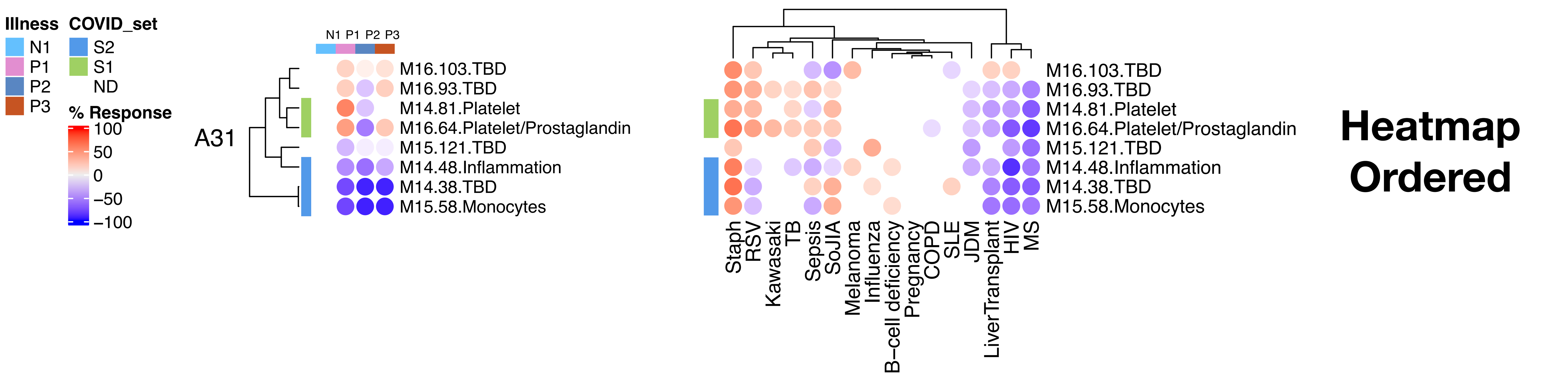

### Aggregate A33: Covid-19 relevant sets S1 & S2

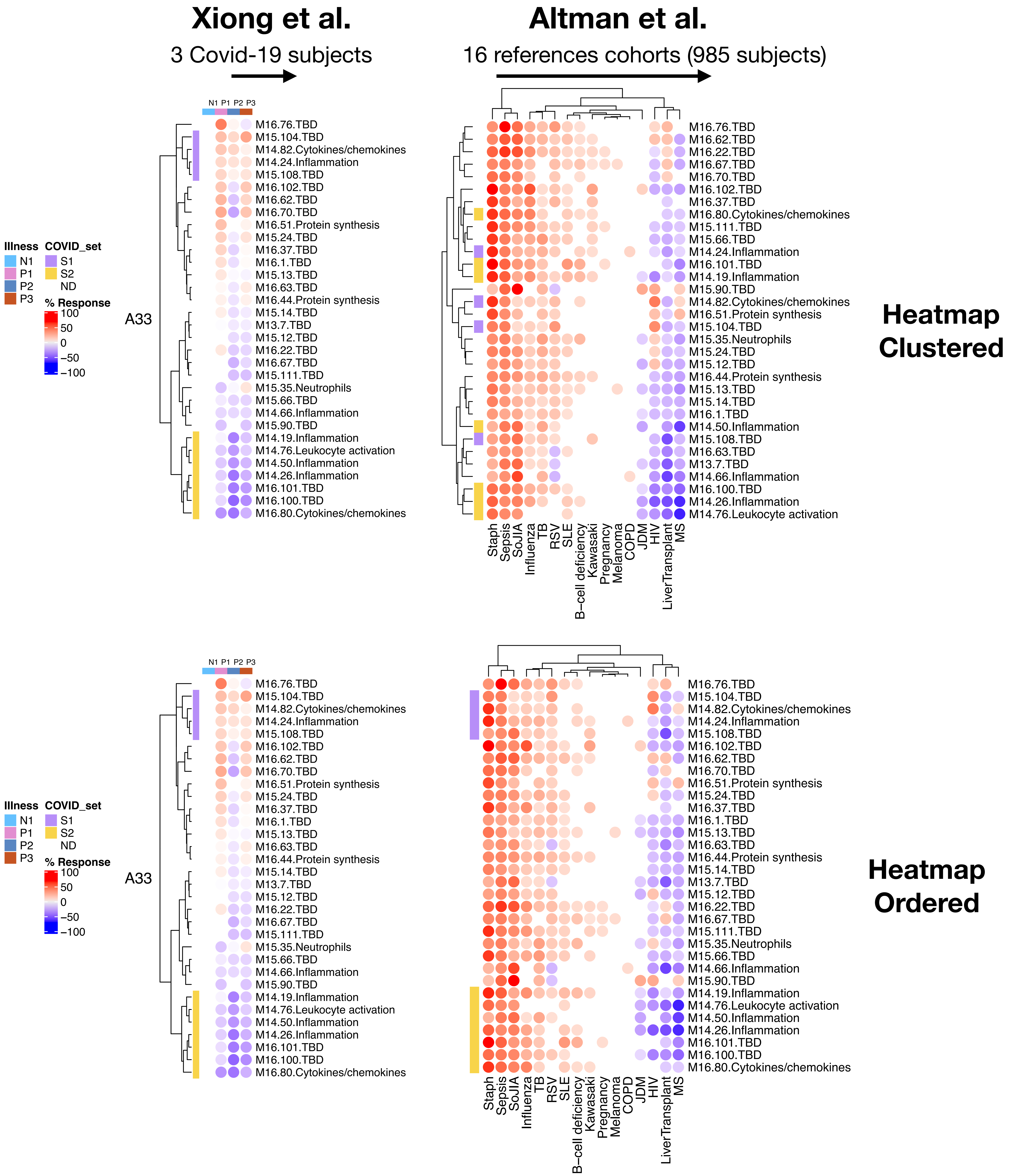

### Aggregate A34: Covid-19 relevant sets S1

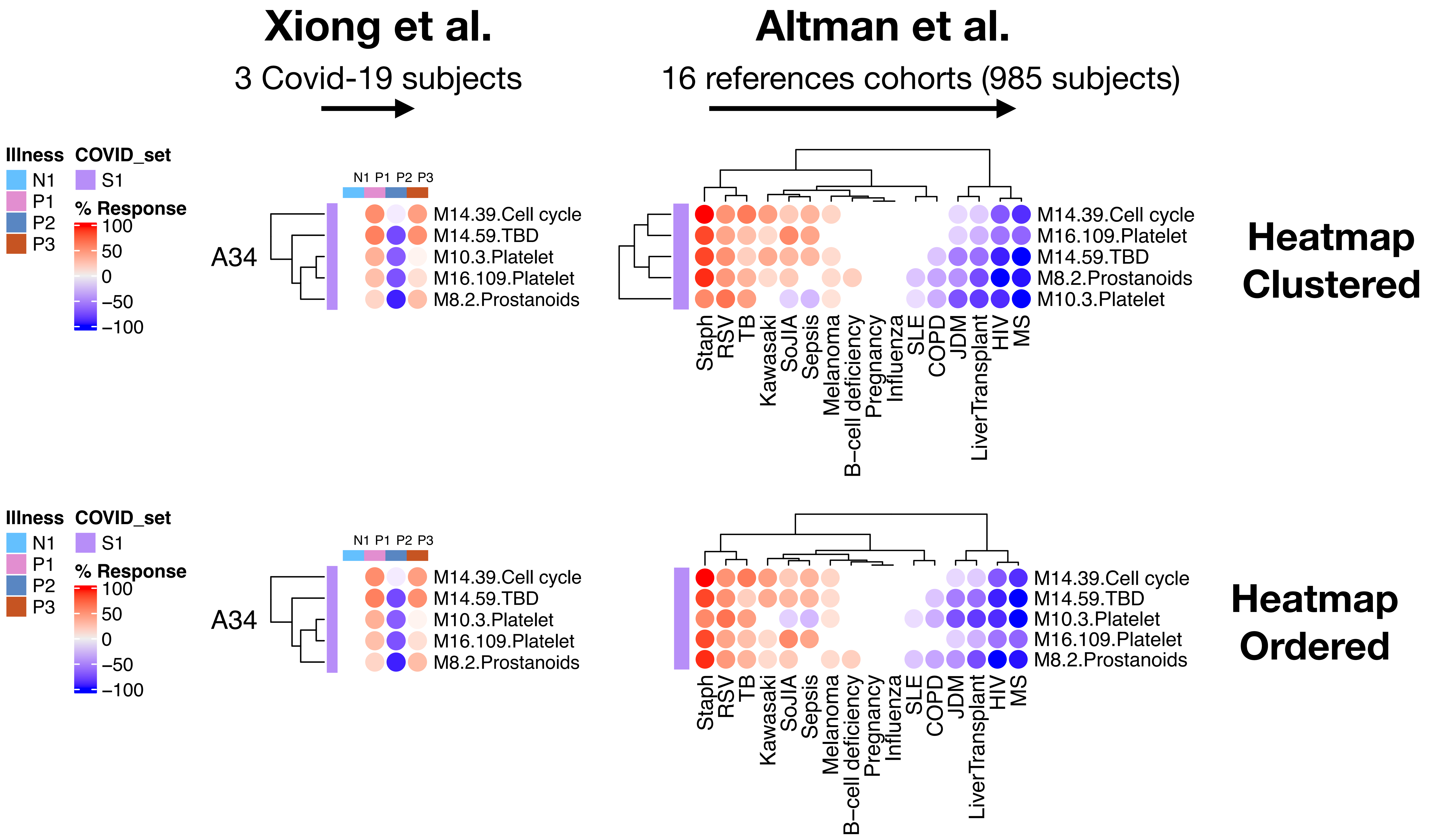

### Aggregate A35: Covid-19 relevant sets S1 & S2

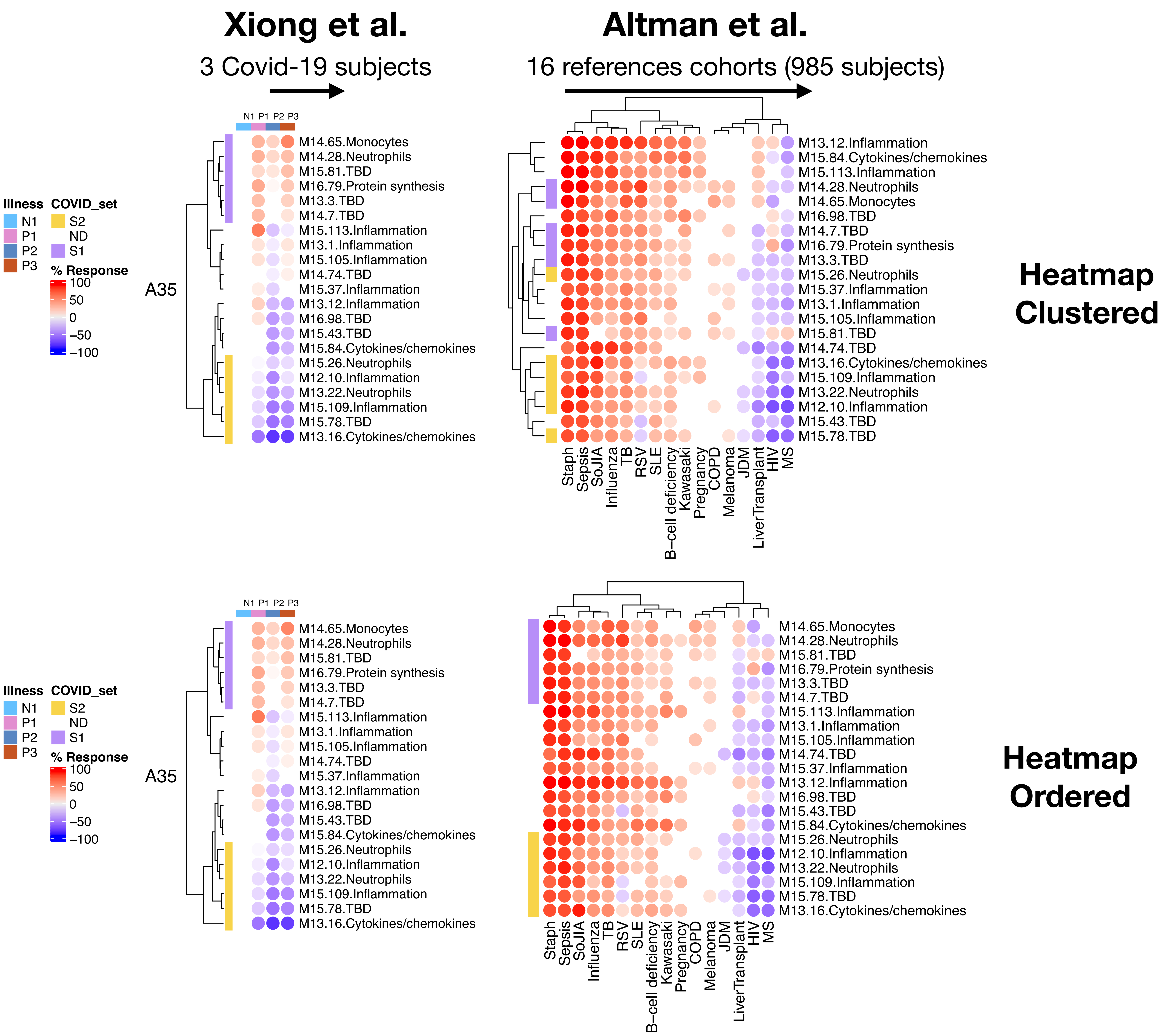

### Aggregate A36: Covid-19 relevant sets S1

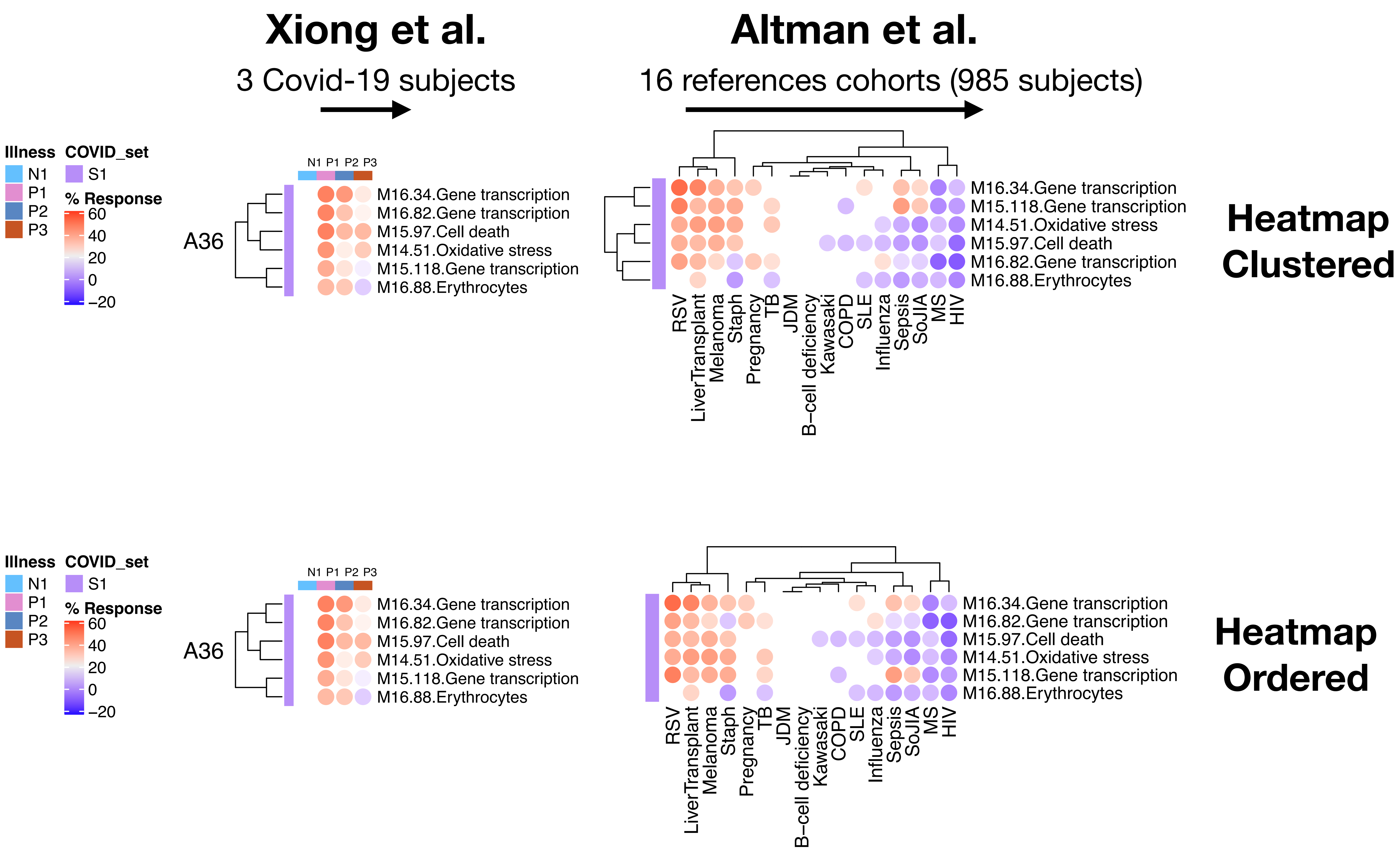

### Aggregate A37: Covid-19 relevant sets S1

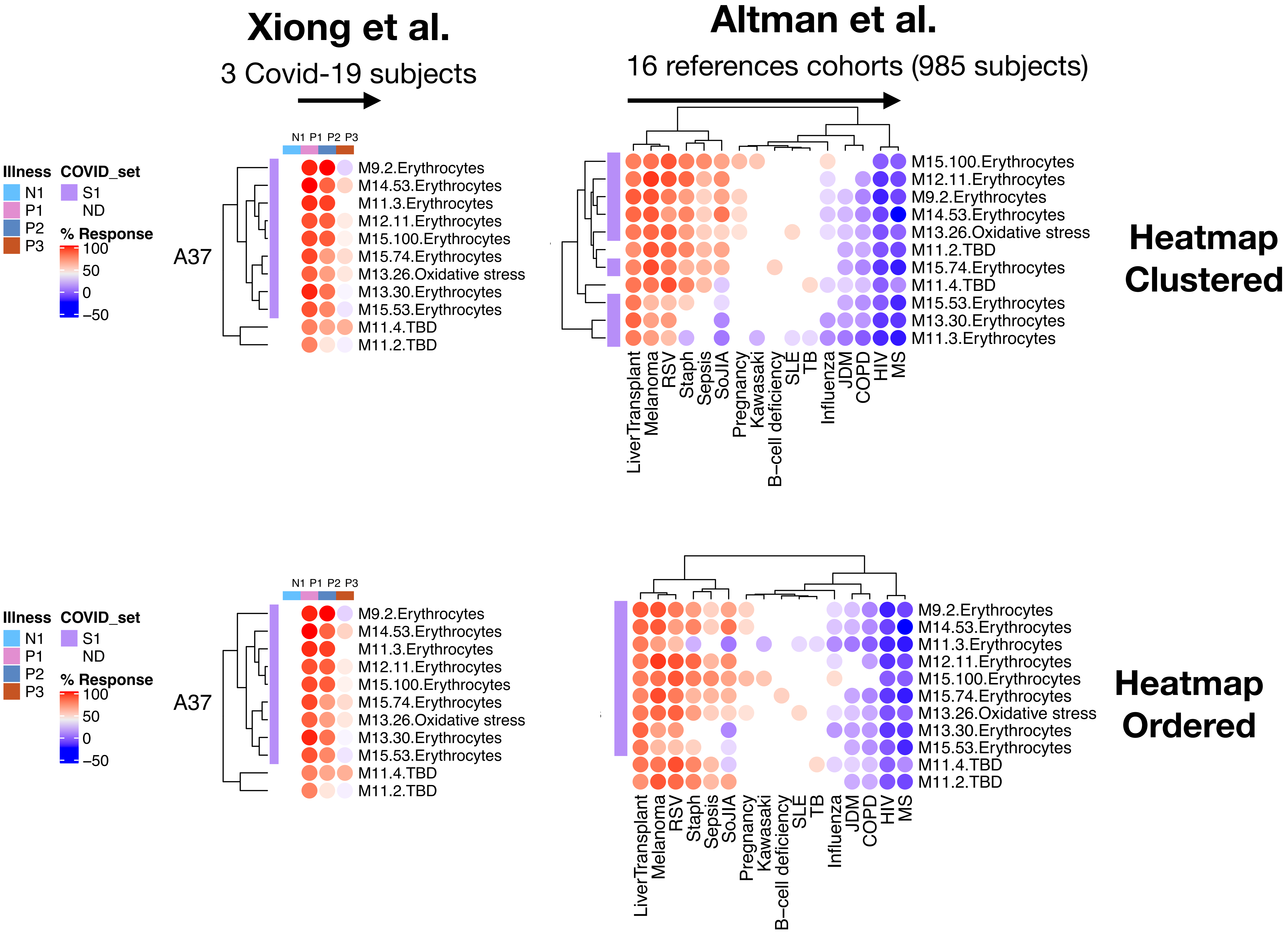

### Aggregate A38: Covid-19 relevant sets S1 & S2

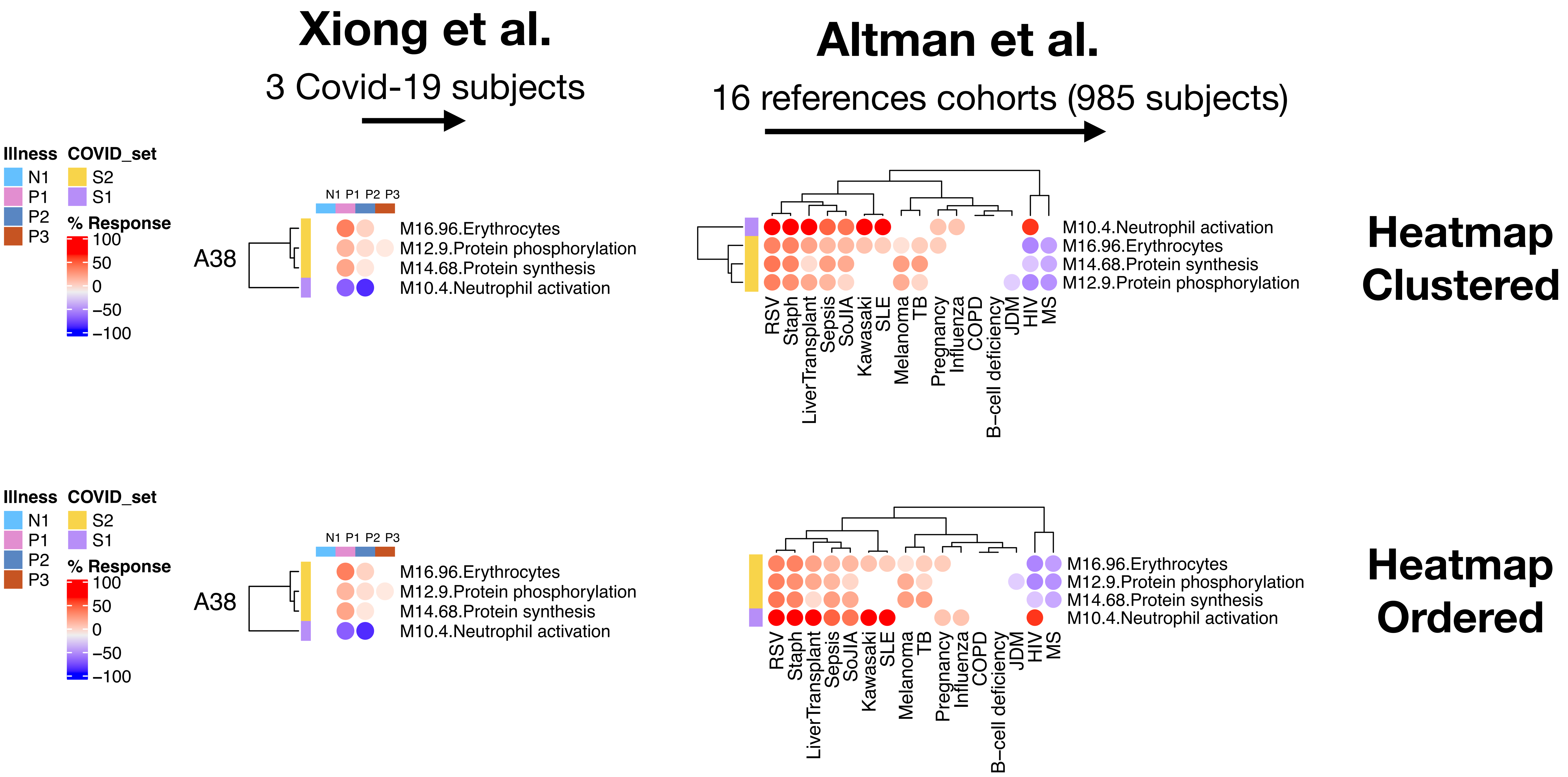
